# Direct observation of conformational dynamics in intrinsically disordered proteins at the single-molecule level

**DOI:** 10.1101/2025.05.26.656162

**Authors:** Saaman Zargarbashi, Cyril Dominguez, Matthew Peters, Arman Yousefi, Sharon Munday, Yanhong Wang, Shreyasi Chatterjee, Andrew Hudson, Reuven Gordon, Christopher J. Mellor, Lei Xu, Mohsen Rahmani, Cuifeng Ying

## Abstract

Intrinsically disordered proteins (IDPs) and intrinsically disordered regions (IDRs) in structured proteins are integral to many biological processes including neurotransmitter regulation, microtubule regulation, and transcription. IDP/IDRs are heterogenous, existing in a conformational ensemble of various interconnected states without a definitive tertiary structure. The high dynamicity of IDPs/IDRs limits ensemble protein characterisation techniques from capturing their properties, and measurements at the single-molecule level are hampered by the necessity to label the protein or modify its microenvironment, affecting their biophysics. Consequently, our understanding of IDPs/IDRs is limited, translating to a lack of knowledge of their roles in related diseases including Alzheimer’s disease, Parkinson’s disease, and various cancers. This work presents the first experimental observation of unmodified IDP/IDR conformational dynamics *in vitro*, at the single-molecule level in real time, achieved by trapping individual IDPs/IDRs in a nanoscale volume using nanoaperture optical tweezers. Our results reveal that IDPs/IDRs exhibit significantly larger conformational variations in solution compared to globular proteins of similar size, as expected. We demonstrate that phosphorylation of native tau-441 by glycogen synthase kinase 3-beta (GSK3β-tau) induces compaction and reduced conformational dynamics. We further observed a disorder-to-order transition during binding of an IDR, the N-terminal region of Src-associated protein in mitosis of 68 kDa (Sam68), to G8.5 RNA. The capability of nanoaperture optical tweezers to monitor the dynamic behaviours of single, unmodified IDPs/IDRs provides a powerful approach to advance our understanding of their elusive behaviours and further decode their roles in associated diseases.

## Main

Intrinsically disordered proteins (IDPs) and intrinsically disordered regions (IDRs) in structured proteins, which constitute approximately 70% of the human proteome^1^, play critical roles in biological processes including neurotransmitter regulation^2^, microtubule regulation^3^, and transcription^4^. Their significant prevalence and roles within various diseases, many of which lack effective treatments and reliable early-stage diagnosis, make IDPs/IDRs important targets for research. Example IDPs include tubulin associated unit (tau) protein and alpha synuclein, which are implicated in neurodegenerative disorders^5^ such as Alzheimer’s disease^6^ and Parkinson’s disease^7^, respectively. IDP/IDR related diseases also extend to cancer^10^ such as the IDR Sam68 (Src-associated protein in mitosis of 68 kDa), which has implicated involvement in ovarian, kidney and lung cancers^11^.

IDPs/IDRs are conformationally heterogenous, dynamically fluctuating between different shapes with variable structure, known as a conformational ensemble^1^. This structural flexibility is often integral to IDP functions, where binding to select targets can induce a particular structure necessary for function, seen in disorder-to-order transitions^12,13^, or retain partial to complete disorder upon binding to form fuzzy complexes^14^. Their heterogeneity and absence of a defined folded state renders many experimental and computational approaches^15^, which were developed for structured proteins, largely inadequate.

Ensemble measurements, such as nuclear magnetic resonance^16^, small angle x-ray scattering^17^ and dynamic light scattering^18^, while very informative, cannot entirely capture the heterogeneity among individual copies of the protein. Structural protein characterisation techniques such as cryogenic electron microscopy (cryo-EM) and x-ray crystallography are suitable for globular proteins, but these techniques yield poor resolution of conformational heterogeneity, leading to low or an absence of electron density in micrographs^19,20^. Additionally, such techniques only capture a snapshot of the protein motion on its conformational landscape, losing dynamic information, which are integral to IDPs/IDRs. Notably, cryo-EM is progressing towards to overcoming some of these limitations, such as the advent of single particle cryo-EM^21^ and use of artificial intelligence to map flexible areas^22^.

Single-molecule fluorescence resonance energy transfer (smFRET) and single-molecule force spectroscopy (smFS) can provide excellent information such as free-energy landscapes^23^, tracking intramolecular displacement^24^, and conformational dynamics^25^, and have even provided insight into IDPs over a decade ago^26,27^. However, the requirement of labelling the protein with an extrinsic fluorophore for smFRET or tethering it to a surface for smFS can perturb the structure and dynamics of the protein^28,29^, particularly for IDPs/IDRs where many are observed to undergo disorder-to-order transitions upon binding^12,13,30^. Currently, no established protein characterisation technique can capture conformational dynamics of label-free IDPs/IDRs at the single-molecule level.

Nanoaperture optical tweezers (NOTs) utilise localised surface plasmon resonance (LSPR) to confine electromagnetic fields to nanoscale volumes (∼tens of nanometres), generating gradient force capable of trapping a single protein molecule a few nanometres in size)^31^. The trapped protein further enhances the electric field through a self-induced back action (SIBA) mechanism, enabling single-protein trapping at low laser power^32^. Additionally, the aperture-based structure effectively dissipates laser-induced heat, making NOTs an ideal platform to study single proteins^33,34^. NOTs have demonstrated their ability in detecting confirmational changes in single molecules in solution without the use of a fluorescent tag or other chemical modifications^35,36^. Through incorporating a microfluidics system to modulate solution conditions whilst the protein is trapped, NOTs have provided information from various processes, including protein-binding interactions^37^, conformational transitions^38^, disassembly kinetics^39^, and energy landscapes^40^.

Here, we use NOTs to monitor conformational fluctuations of label-free IDPs/IDRs in solution, elucidate their free-energy landscapes, and observe a disorder-to-order transition of an IDR upon binding to RNA. We demonstrate this with a focus on three different IDPs/IDRs associated with disease progression: native tau-441, tau-441 phosphorylated by glycogen synthase kinase 3-beta (GSK3β-tau) and the N-terminal region of Sam68. This work reveals differences in structural flexibility between single IDPs and globular proteins of similar size, provides experimental evidence of structural changes due to phosphorylation of IDPs, and the trajectory of a single IDR transitioning between disordered and ordered structures upon binding and unbinding of nucleic acid binding partners, with well resolved binding kinetics. All the above insights into the conformational dynamics of label-free IDPs/IDRs were previously inaccessible to any single-molecule approach.

### Transmission traces reflect protein conformational dynamics

Trapping of either an IDP/IDR or a globular protein is achieved using a gold double-nanohole (DNH) structure (Fig. 1a; full setup in Fig. S1). An 852 nm laser excites the LSPR of the DNH, generating strong field enhancement at the DNH gap (Fig. S2a) to trap a single protein molecule. In this work, we trap all proteins with the laser power that produces a local temperature of ∼37°C (i.e., 20 mW, see Figs. S3a and S3b). When a molecule enters the trapping region, the disparity in the refractive index between the trapped molecule and the surrounding media affects the light scattered by the nanoaperture (Figs. S2b and S2c). This scattering directly correlates to the polarisability of the trapped protein, which is determined by its volume (linear relation, Fig. S2d), conformation and orientation (Fig. S2e), and dielectric constant^41,42^. The forward-scattered light is collected by an objective (NA = 0.1) and then recorded as the transmission intensity (*I*) by an avalanche photodiode (APD)^43^ (Fig. S1). We analyse the protein dynamics using the normalised transmission intensity change, Δ*I/I*_*0*_, where *I*_*0*_ is the baseline intensity of an unoccupied DNH, and Δ*I* = *I* - *I*_*0*_ represents the change in transmission intensity from this baseline (Fig. 1b and Fig. S1).

**Fig. 1:**
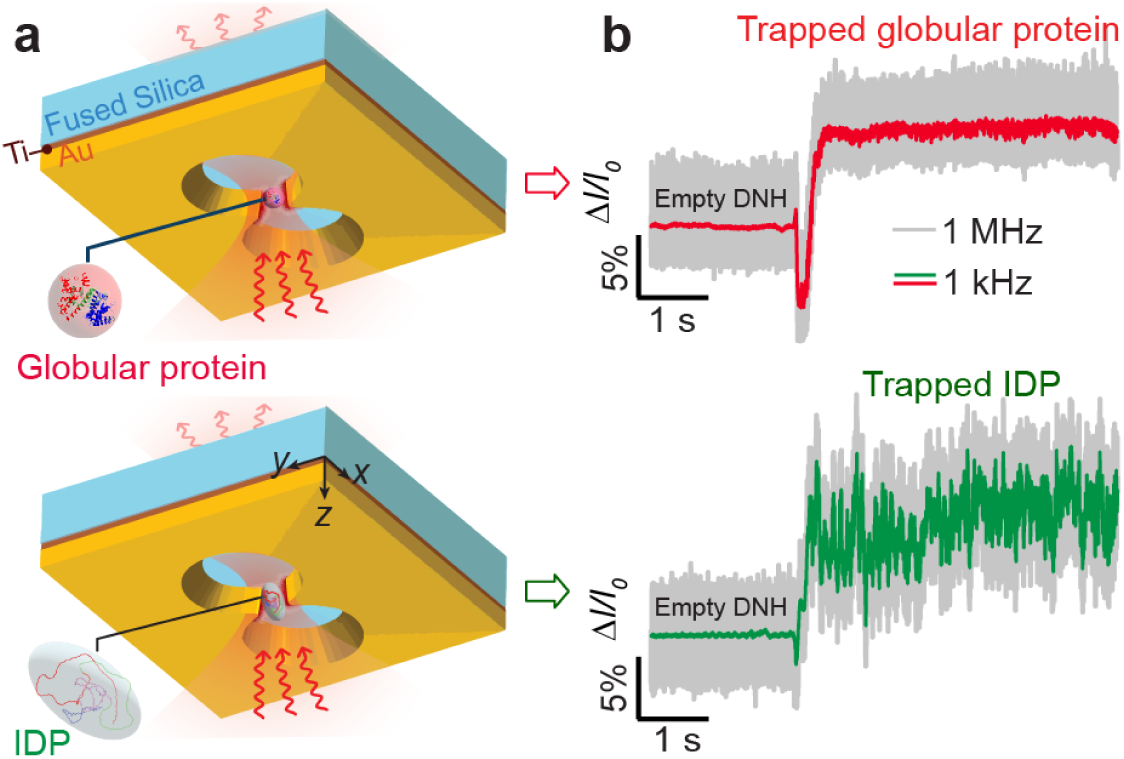
Single-molecule protein trapping using nanoaperture optical tweezers. **a**, Schematic of label-free trapping of either a globular protein (top) or an IDP (bottom) within a DNH structure. The DNH was fabricated in a 100-nm gold film and passivated with PEG-thiol (see materials and methods for detailed parameters). SEM images of representative DNH geometries are shown in Fig. S4. **b**, Representative trapping traces for a globular protein (BSA) and an IDP (GSK3β-tau). Plotted trace consists of raw data collected at 1 MHz sampling rate (light grey) and data filtered at 1 kHz (globular protein: red, IDP: green). Greater variations in the normalised transmission intensity (Δ*I/I0*) indicate greater conformational dynamics. IDP and globular protein structures were generated using AlphaFold 3^47^ and edited with ChimeraX^48^.

Comparing two representative trapping trajectories for an IDP and a globular protein reveals differences in their properties (Fig. 1b). The initial increase in Δ*I/I*_*0*_ indicates the protein entering the DNH, with the magnitude linearly scaling with protein size (Fig. S2d). While the globular protein maintains relatively low Δ*I/I*_*0*_ fluctuations, consistent with its rigid structure, the IDP trace exhibits significantly larger signal variations. Two factors may contribute to this difference. First, the loose, extended conformation of IDPs exposes a greater surface area to the solvent, resulting in a larger hydration shell with higher water density than that of globular proteins^44,45^. This hydration effect increases the local refractive index around the protein in its elongated states, leading to greater transmission changes. Second, elongated particles of equivalent volume produce orientation-dependent Δ*I/I*_*0*_ signals, as demonstrated by our simulations (Fig. S2e). IDPs/IDRs continuously switch between different extended conformations on microsecond-to-second timescales^46^, producing intrinsic Δ*I/I*_*0*_ fluctuations that reflect their conformational sampling and orientations. We hypothesise that elongated globular proteins may bias in a preferred trapping orientation that maximises the local electric field through the SIBA mechanism, and consequently, results in a higher Δ*I/I*_*0*_ value than their spherical counterparts, as previously reported^31,37,40^. As detailed in Supplementary SI-2 and SI-3, Δ*I*/*I*_*0*_ depends critically on the refractive index, shape, and orientation of the protein within the DNH gap. These dependencies enable Δ*I*/*I*_*0*_ to serve as an indicator for the global compactness of the trapped protein.

### Experimental insights into ordered and disordered proteins

Figures 2a-2c compare the transmission-time traces corresponding to the trapping of a native tau-441 and its phosphorylated variant GSK3β-tau, with those of the globular protein haemoglobin. Trapping these proteins using DNHs with similar dimensions resulted in comparable Δ*I*/*I*_*0*_ of approximately 0.1 (Figs. 2a–c), due to their similar molecular weight – haemoglobin, 64.5 kDa, native tau-441, 45.9 kDa, and GSK3β-tau, ∼46–48 kDa. The extended conformations of IDPs result in higher polarisability than globular proteins of similar molecular weight, which explains why native tau-441 and GSK3β-tau exhibit Δ*I/I*_*0*_ values comparable to haemoglobin despite their lower molecular weight. The trace segments in Fig. 2d reveal substantially different fluctuations in Δ*I/I*_*0*_ between the three proteins. Native tau-441 and GSK3β-tau resulted in larger fluctuations compared to haemoglobin (Fig. 2d inset), consistent with the higher flexibility of IDPs relative to globular proteins. This trend is observed across multiple trapping traces involving various IDPs/IDRs and globular proteins (Figs. S5a and S5b) rather than specific to tau and haemoglobin. Figure S5c confirms that IDPs/IDRs demonstrate higher normalised root-mean-square (NRMS), and therefore dynamics, than globular proteins of similar molecular weight. When the laser is turned off for several seconds then turned back on, the transmission intensity returns to the baseline level (Fig. 2a), indicating that the protein molecule diffused away from the trap without surface adsorption. These results demonstrate the effectiveness of the PEG-thiol coating in minimising nonspecific protein adsorption—a major challenge when studying IDPs/IDRs, particularly under conditions where the trapping force retains them close to the gold surface for extended periods. Occasionally, however, the protein did not diffuse away after the laser was turned off, as shown in Fig. S6. These events were excluded from subsequent analysis and discussion in this work.

**Fig. 2:**
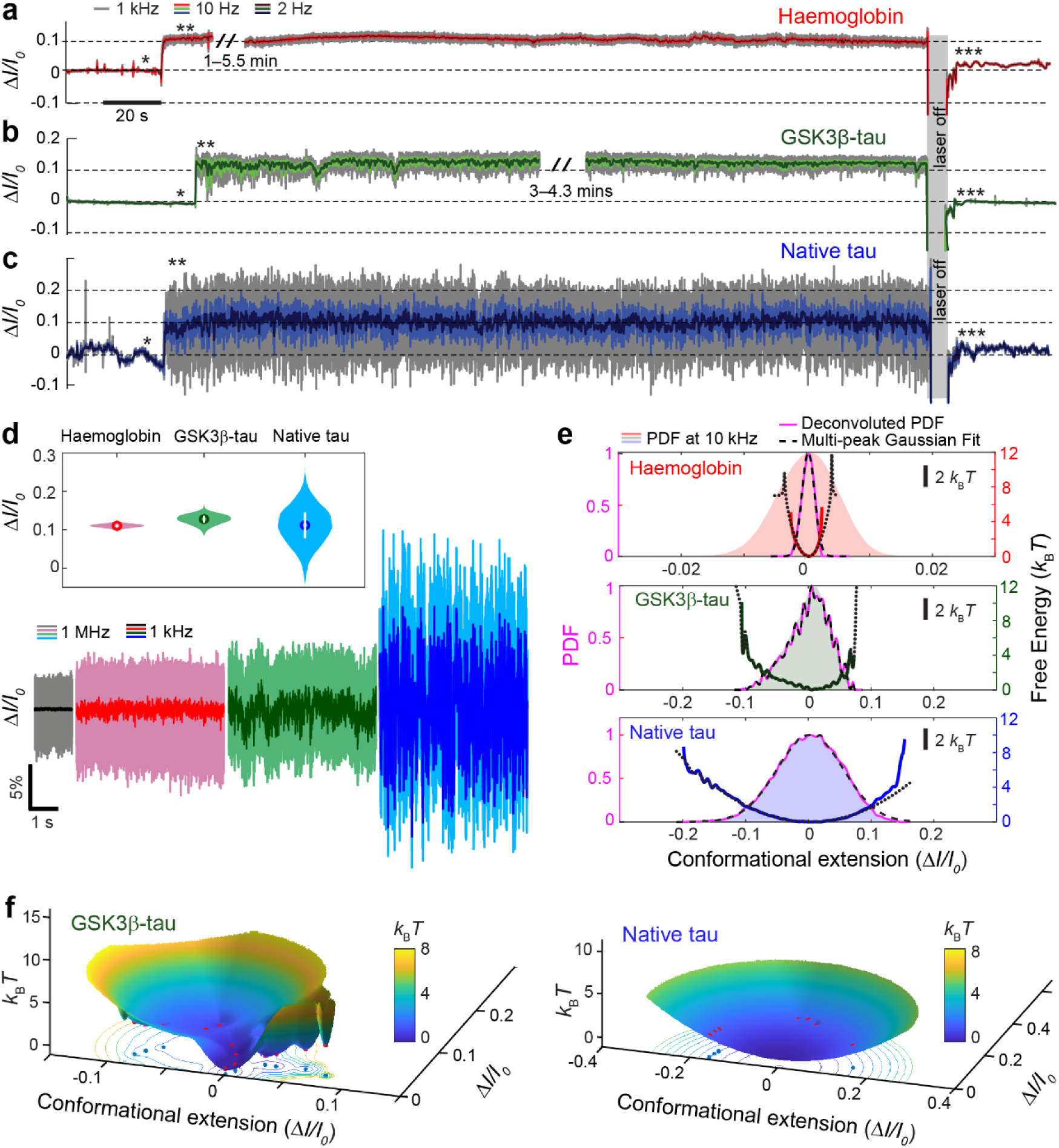
Revealing protein dynamics and energy landscapes using nanoaperture optical tweezers. **a–c**, Optical transmission traces over 6 minutes through a double nanohole (DNH) for trapped proteins: **a**, haemoglobin (64.5 kDa); **b**, GSK3β-tau (∼46–48 kDa); and **c**, native tau-441 (45.9 kDa). Transmission intensities were sampled at 1 MHz and digitally filtered at 1 kHz (grey), 10 Hz (light coloured) and 2 Hz (dark coloured). Raw traces are shown in Fig. S7. Asterisks indicate the following events: baseline before trapping (*), protein trapped (**), and protein released (***). **d**, Zoomed segments taken from **a–c**, comparing baseline (black), haemoglobin (red), GSK3β-tau (green), and native tau-441 (blue). Data are shown in raw (1 MHz) and filtered at 1 kHz. Inset: violin plots of Δ*I/I*_*0*_ values during protein trapping. **e**, Probability density functions (PDFs) of Δ*I/I*_*0*_ over 20 s of 10 kHz digital filtered data for each protein (colour shaded regions), with deconvoluted PDFs (magenta curves) fitted using multi-peak Gaussian models (black dashed lines). 2D energy landscapes were derived from the deconvoluted PDFs for haemoglobin (red line), GSK3β-tau (green line), and native tau-441 (blue line). **f**, 3D energy landscapes over 20 s for GSK3β-tau and native tau-441, constructed from the deconvoluted PDFs in **e**. Distance from the origin (0, 0, 0) reflects conformational extension (Δ*I/I*_*0*_ magnitude, *x*- and *y-*axis) and thermodynamic stability (*k*_B_*T, z-*axis). All data were acquired at 37°C. See Supplementary SI-9 for details on PDF deconvolution and the calculation of energy landscapes.

Probability density functions (PDFs) of the transmitted intensity reveal how frequently a protein samples different conformations, providing valuable insight into its accessible structural states. Deconvolution of the PDFs using a point spread function removes the translational and rotational motion of proteins and allows extraction of the PDFs associated with the true protein conformational changes. Figure 2e shows that the deconvoluted PDF for haemoglobin reveals a single, sharp peak, consistent with it existing predominantly in a single conformational state, characteristic of a globular protein. In contrast, the deconvoluted PDFs of both GSK3β-tau and native tau-441 exhibit broader Δ*I/I*_*0*_ distributions, indicating greater conformational variability and a more dynamic behaviour than haemoglobin. Native tau-441 displays the broadest distribution, suggesting it samples a wider conformational landscape and possess greater structural flexibility than GSK3β-tau.

Converting the PDF of single-molecule folding trajectories into free-energy landscapes is well established in techniques such as FRET^49^ and smFS^50^. These landscapes provide valuable insights into protein folding, including the number of distinct conformational states, their relative free-energy differences, and how these are altered by binding interactions^51^. Recently, NOTs have been used to resolve the energy landscape of a single unmodified protein^40^. For label-free monomeric IDPs/IDRs, experimental derivation of energy landscapes is particularly important for understanding their conformational dynamics, molecular interactions, and biological functions^51^. Until now, this information has been accessible only through computational modelling^52,53^, due to the challenges of experimentally measuring label-free IDPs/IDRs at the single-molecule level.

Here, we present the first experimental measurement of the free-energy landscape of label-free IDPs at the single-molecule level. 2D energy landscapes were calculated by taking the negative logarithm of the deconvoluted PDFs, as described by Eq. S3, and are shown in Fig. 2e (red, blue and green curves). Relative free-energy values are expressed in units of *k*_B_*T*, where *k*_B_ is Boltzmann’s constant and *T* is temperature, representing the energy available to the protein. Lower *k*_B_*T* values correspond to more stable conformational states. As expected, Figure 2e reveals that haemoglobin exhibits a sharp funnel-like landscape characteristic of globular proteins^54^, whereas native tau-441 and GSK3β-tau display multiple shallow minima with lower energy barriers between them, consistent with a broad ensemble of conformational states typical of IDPs^54^. To identify predominant conformational states, we applied multi-peak Gaussian fits to the deconvoluted PDFs, (black dashed curves in Fig. 2e, with individual peaks presented in Fig. S8a). We then used these fitted PDFs to generate fitted energy landscapes (black dotted curves, Fig. 2e). It is worth noting that we deliberately overfit the PDFs to maximise accuracy in reconstructing the free-energy landscapes (Figs. S8a and S8b), resulting in fitted energy landscapes that closely follow the measured curve.

To better visualise the complex conformational dynamics of these IDPs, we converted the 2D energy landscapes into 3D plots by mapping peak positions onto a polar coordinate system (Fig. S9). In these plots, angular coordinates reflect the relative spatial distribution of conformational states, while peak amplitudes correspond to the negative logarithm of the fitted PDF—such that higher probability indicates lower free energy. Haemoglobin, as expected for a globular protein, displays a single, funnel-shaped energy landscape consistent with a single, stable folded conformation (Fig. S10). For native tau-441, the 3D energy landscapes reveal multiple shallow troughs with low energy barriers, consistent with the conformational heterogeneity expected in IDPs (Fig. 2f). In contrast, alongside shallow troughs similar to those observed in native tau-441, GSK3β-tau exhibits several deep, well-defined troughs, indicating the presence of distinct and stable conformational states (Fig. 2f). These states are also apparent in the top-down projection shown in Fig. S11. The continuous energy landscapes over 120-seconds from native tau-441 and GSK3β-tau (Fig. S12) confirm conformational dynamics consistent to those observed in the 20 s intervals (Fig. S12).

### Phosphorylation induced order of tau by GSK3β

The results shown in Fig. 2 and Fig. S11 suggest that GSK3β phosphorylation of native tau-441 induces increased order and compaction. Two mechanisms likely underlie these changes: electrostatic interactions and secondary structure formation. First, native tau-441 has a theoretical isoelectric point of ∼8.24, carrying a net positive charge at pH 7.2. Phosphorylation introduces negatively charged phosphate groups, subsequently reducing the net charge and promoting compaction, consistent with previous reports^55,56^. Second, phosphorylation may shift local secondary structure. GSK3β phosphorylation has been shown to increase α-helix propensity at the expense of polyproline type II (PPII) helices within the proline-rich domain of tau^57^ (Fig. S13). As α-helices are shorter than PPII helices (5.4 Å/turn vs. 9.3 Å/turn)^58^, this structural transition would also induce compaction in native tau-441. Supplementary SI-10 provides an in-depth discussion of potential phosphorylation effects.

To confirm that GSK3β phosphorylation shifts native tau-441 towards more compact and ordered conformations, we performed additional trapping experiments shown in Figure 3. Dataset 1 was acquired using a single DNH structure, while Dataset 2 used two similarly sized DNHs on the same chip (see Fig. S5 for DNH size variation). GSK3β-tau again exhibited reduced dynamic behaviour compared to native tau-441 (Figs. 3a and 3b). Deconvoluted PDFs and energy landscapes show more compact conformations in GSK3β-tau, as indicated by smaller Δ*I/I*_*0*_ fluctuations (left panels, Figs. 3c and 3d). To further investigate the timescale of conformational dynamics, we analysed the power spectral density (PSD) of each dataset (right panels, Figs. 3c and 3d). Compared to native tau-441, GSK3β-tau displayed lower power fluctuations in the 1 Hz–1 kHz range, indicating its increased order and restricted large-scale dynamics occurring on the millisecond–second timescale. We note that this platform resolves complete conformational transitions on ∼10 ms timescales for both tau-441 and GSK3β-tau, as estimated by cross-mean frequency analysis of transmission traces (Fig. S14).

**Fig. 3:**
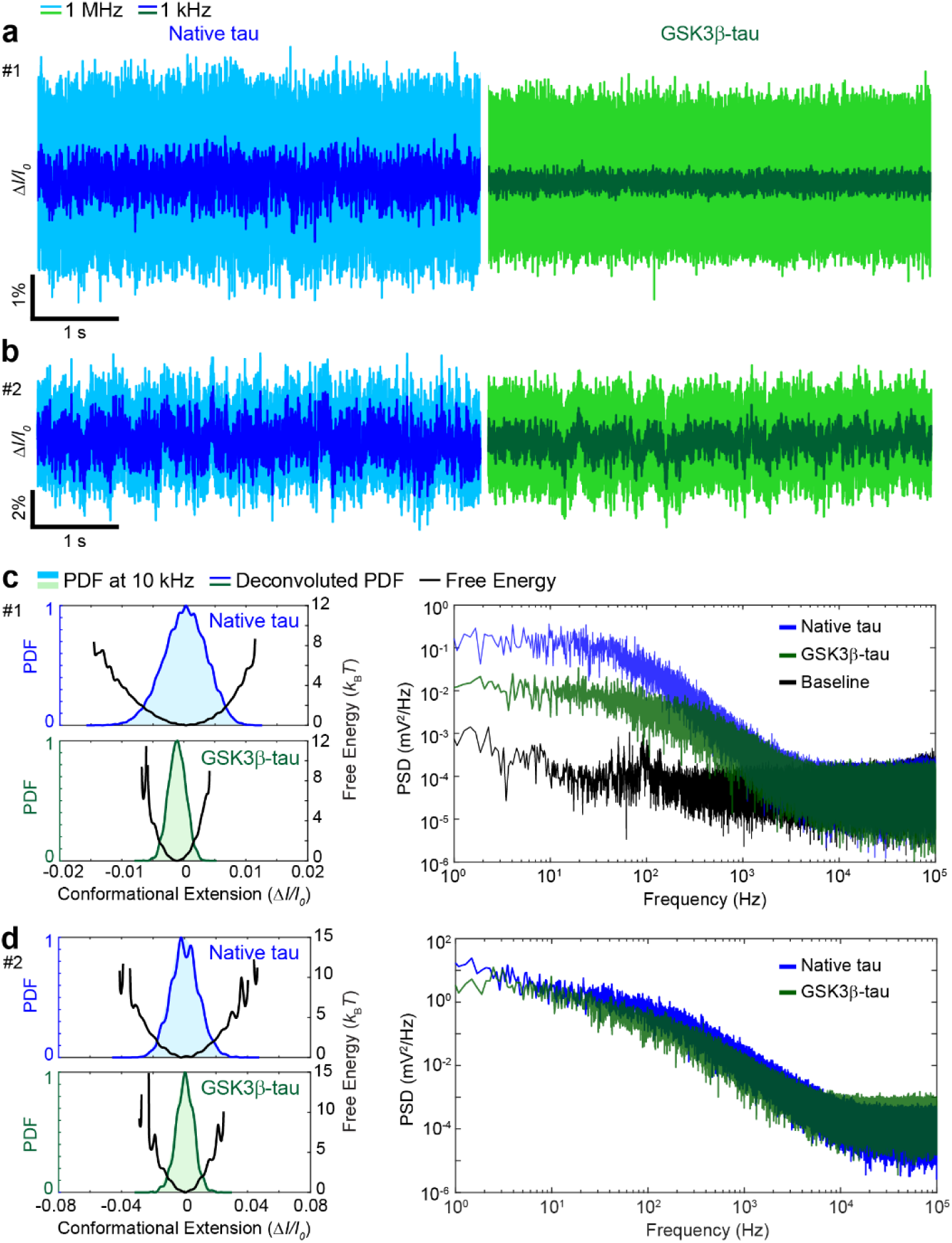
Effect of GSK3β phosphorylation on tau-441. **a, b**, Five-second transmission traces comparing native tau (blue) and GSK3β-tau (green), shown as both raw (1 MHz, light blue or light green) and filtered (1 kHz, dark blue or dark green). **c, d**, Left: PDFs of 20-s traces (10 kHz filtered, light blue or green shades) with deconvoluted PDFs (dark blue or green), along with 2D energy landscapes (black) for datasets in **a** and **b** respectively. Right: corresponding PSD plots for the 20-s traces. Dataset #1 was obtained using a third-party fabricated DNH with a reduced gap size (see red box in Fig. S4 for SEM images).

### Disorder to order transition of the Sam68 N-terminal

Our results demonstrate that NOTs enable the direct and prolonged observation of conformational dynamics in single, unmodified IDPs, and allow reconstruction of their free-energy landscapes. To illustrate the broader utility of this approach in capturing disorder-to-order transitions at the single-molecule level, we investigated the RNA binding kinetics of Sam68—a system not previously studied using single-molecule methods.

Sam68 is an RNA-binding protein composed of three main regions: the N-terminal (residues 1– 96), the central STAR binding domain (residues 97–260), and the C-terminal (residues 261–443)_59_. It binds to G8.5 RNA—a 40-nt RNA sequence—with a dissociation constant (*K*_d_) of ∼12 nM determined from an electrophoretic mobility shift assay^60^. This affinity, however, is significantly reduced to around 36.1 µM when only the central STAR domain is present^61^, suggesting that the intrinsically disordered N- and C-terminal regions contribute to strong binding. NMR experiments later confirmed that both termini independently bind G8.5 RNA, with affinities of 1–10 µM for the N-terminus and 30–70 µM for the C-terminus^62^.

Here, we focus on the Sam68 N-terminal IDR and its interaction with G8.5 RNA, where the RNA-binding activity is attributed to the two arginine/glycine (R/G)-rich motifs—^45^RGGGGG^50^ and ^52^RGG^54^ on the Sam68 N-terminal—and the adenosine/uracil (A/U) rich motif ^22^AUUAAAA^28^ in G8.5 RNA^63^ (full sequences are presented in Figs. S15a and S15b). We trapped the N-terminal of Sam68 for ∼20 minutes before introducing 1 µM G8.5 RNA to monitor binding dynamics (Fig. 4a). Upon RNA arrival at the trapping site (∼30 min after initial trapping), a sharp increase in transmission occurs, indicative of a higher polarisability for the RNA-bound complex compared to the unbound protein. This increase is accompanied by a significant reduction in signal fluctuations (Figs. 4a and 4b), consistent with a transition to a more stable and ordered structure, representing the first direct observation of a disorder-to-order transition in the Sam68 N-terminal IDR.

**Fig. 4:**
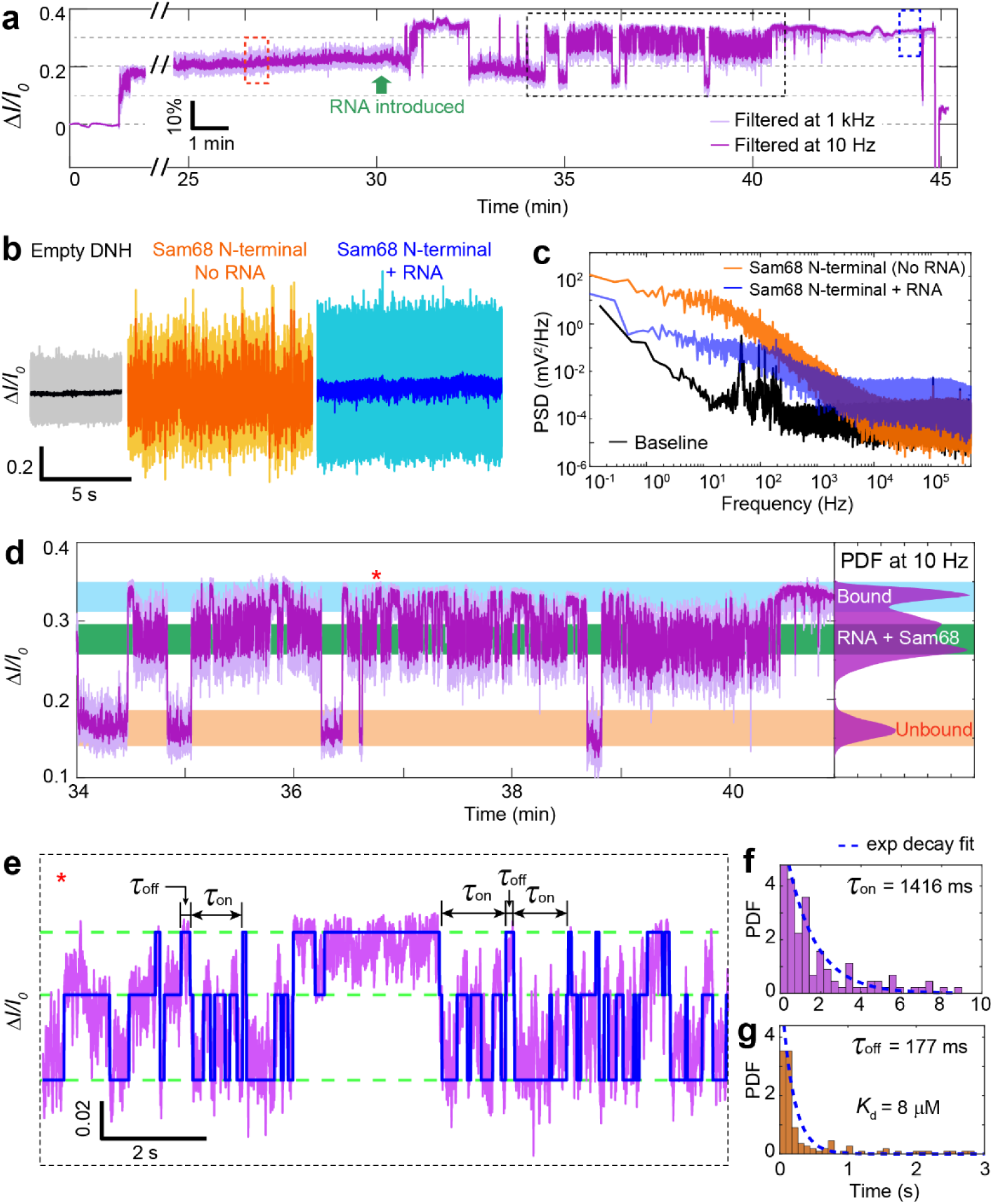
Binding of G8.5 RNA to the Sam68 N-terminal. **a**, Transmission trace of optical trapping of the Sam68 N-terminal followed by binding/unbinding of RNA. Data was filtered to 1 kHz (light purple) and 10 Hz (dark purple). Raw trace available in Fig. S16. **b**, Zoomed segments from **a** depicting the baseline, Sam68 N-terminal before RNA binding (orange, from orange dashed box) and after RNA binding (blue, from blue dashed box). Traces show raw data (1 MHz, light colours) and filtered data (1 kHz, dark colours). **c**, PSD of Sam68 before RNA binding (orange), after RNA binding (blue) and baseline (black) for comparison. **d**, 7-minute segment from **a** (black dashed box) depicting conformational fluctuations of the Sam68 N-terminal between the unbound (orange) and bound (blue) states to RNA, in addition to a condition in which both components are unassociated in the optical trap (green). Right inset: PDFs for the 7-min trace filtered at 10 Hz. **e**, Eight-second zoomed trace from **d** (red asterisk) using a 3-step fit to assign levels to the RNA-bound, and two intermediate states for cohabitation of the unbound Sam68 N-terminal and RNA. τ_on_ and τ_off_ were assigned for binding time and dissociation time, respectively. **f, g**, Histograms of τ_on_ and τ_off_ obtained from the whole trace in **d**, overlaid with single exponential decay fits (blue dashed curves). The dissociation constant (*K*_d_) for the binding interaction were calculated using the average τ_on_ and τ_off_ with details provided in Supplementary SI-14.

Figure 4c compares PSDs of the RNA-bound and unbound states of N-terminal region. The RNA-bound state exhibits reduced signal fluctuations compared to the unbound state, particularly at frequencies below 1 kHz (>1 ms), suggesting restricted dynamics and enhanced structural stability induced by RNA binding. This binding is reversible, with the signal returning to pre-binding levels upon RNA dissociation. The transmission trajectory during binding/unbinding shows three discrete transmission levels (Fig. 4d), with PDF peaks corresponding to: (i) the unbound Sam68 N-terminal, (ii) the RNA-bound state, and (iii) an intermediate level likely reflecting co-localisation of Sam68 and G8.5 RNA within the trap without complex formation. To quantify these binding kinetics, we applied a three-step fitting model, assigning levels to the bound and unbound states (Fig. 4e). Exponential decay fits applied to the histogram of association time (τ_on_) and dissociation time (τ_off_) yielded average τ_on_ and τ_off_ values of 1416 ms and 177 ms, respectively (Figs. 4f and 4g). From these values, we estimated a dissociate constant *K*_d_ of ∼8 µM, consistent with previous NMR results (1-10 μM)^62^. Details of *K*_d_ calculation are available in Supplementary SI-14. Notably, in some instances, RNA binding induced a transition to a persistent ordered conformation (Fig. S17), likely due to the relatively high affinity. While such traces cannot quantify the dissociation constant, they consistently showed attenuated signal fluctuations in the RNA-bound state, with dynamics reduced on timescales of hundreds of microseconds to seconds. Additional context for this observation is provided in Supplementary SI-15.

## Conclusions

This work establishes nanoaperture optical tweezers (NOTs) as a powerful platform for directly probing the conformational dynamics of single, label-free intrinsically disordered proteins (IDPs) and intrinsically disordered regions in structured proteins (IDRs) in solution. The optical trapping signals reveal how IDP/IDR structural disorder manifests through distinct intensity fluctuations, energy states, and dynamic timescales, compared to globular proteins. At single-molecule resolution, we provide experimental evidence that phosphorylation of native tau-441 by GSK3β reduces conformational heterogeneity, with suppressed dynamics observed on the millisecond– second timescale. We further demonstrate the real-time observation of a disorder-to-order transition in the Sam68 N-terminal region upon RNA binding, with dissociation kinetics consistent with ensemble measurements.

These findings highlight the capacity of NOTs to resolve binding-induced structural transitions and conformational kinetics at the single-molecule level, while circumventing key limitations of existing single-molecule techniques. Unlike fluorescence-based methods, this label-free approach avoids photobleaching, background fluorescence interference, and the challenge of distinguishing fluorophore dynamics from protein motions. Compared to single-molecule force spectroscopy, it eliminates the need for specific tethering, which often requires prior structural knowledge of the protein.

Nevertheless, the approach has limitations. It lacks of spatial resolution for monitoring distance changes between specific residues or identifying precise interaction sites. Instead, it captures how conformational rearrangements alter the local refractive index within the optical hotspot, offering a complementary perspective to structure-specific single-molecule techniques. Signal variability between individual DNH structures, due to inevitable nanofabrication imperfections, further complicates quantitative comparisons across traps from different structures. Optical trapping forces may also perturb protein conformations, though this effect remains to be fully characterised. In terms of temporal resolution, while the avalanche photodiode offers 50 MHz bandwidth, the observable protein dynamics are constrained within ∼10 kHz by the background noise from empty DNH and laser sources. Future improvements may focus on improving temporal resolution, for example by detecting large-angle scattering to reduce background noise, or by employing cross-correlation between two detection angles to counteract laser source fluctuations. Integration with complementary techniques, such as nanopore sensing and/or Raman spectroscopy, could enhance trapping efficiency while providing orthogonal single-molecule validation. To mitigate signal variations across DNHs, the incorporation of microfluidic systems for extended-duration protein trapping, similar to the Sam68 RNA-binding experiment presented here, would enable systematic observation of conformational responses to controlled physicochemical changes (e.g., pH, ionic strength, or ligand concentration) without requiring repeated trapping events.

Taken together, the approach presented here provides unique insights into the conformational dynamics of disordered proteins that are inaccessible to conventional single-molecule techniques. These findings position NOTs as a versatile and minimally invasive technique for investigating IDP conformational dynamics, molecular interactions, and post-translational modifications, expanding the current experimental toolkit for studying protein disorder and its role in biological function and disease.

## Acknowledgements

This research work was supported by the UK-India Education Research Initiative (UKIERI) and the Academy of Medical Sciences (AMS) Springboard Award. S.Z. acknowledges support from the Biotechnology and Biological Sciences Research Council Doctoral Training Partnerships (BBSRC DTP) (BB/T0083690/1). M.R. appreciates the support from the Royal Society and the Wolfson Foundation. C.D and A.H. acknowledge support from the BBSRC sLoLa grant (BB/T000627/1).

## Author contributions

C.Y. conceived the project. S.Z. and Y.W. performed trapping experiments for all proteins and prepared all non-Sam68 protein buffers. S.M. prepared Sam68 constructs. S.Z. performed nanofabrication of DNHs. A.Y. provided support for trapping experiments and nanofabrication. C.D and A.H. provided support for Sam68 content. C.Y. and M.P. conducted energy landscape calculations. S.Z. and C.Y. performed the data analysis. S.C. provided guidance on native tau-441 and GSK3β-tau. R.G. provided guidance on plasmonic nanotweezers and data interpretation. C.J.M., L.X., M.R., and C.Y. provided S.Z. supervisory guidance. The manuscript was written by S.Z., with assistance from C.D., M.P., A.Y., S.C., A.H., R.G., C.J.M., L.X., M.R., and C.Y.

## Materials and Methods

### Fabrication of double nanohole structures in gold film

The double nanohole (DNH) structures used in this work were fabricated as previously described^1–3^. A 550 μm thick fused silica wafer was coated with a 30 nm silicon nitride layer using low-pressure chemical vapor deposition (LPCVD). Subsequently, a 5 nm titanium layer and a 100 nm gold layer were deposited using electron-beam evaporation at 190 °C. The wafers were diced to 10 mm × 10 mm chips for further use. Two nanoholes, each with a depth of 90 nm, diameter of 160 nm and a centre-to-centre distance of 200 nm were etched into the gold layer using a focused ion beam (FIB, Zeiss Crossbeam) with a gallium ion source. A rectangle of 3 nm in height was then etched to connect the edges of the two holes to form the DNH gap. The FIB was operated at 30 kV with a beam current of 1 pA.

### Double nanohole surface passivation

Double nanohole structures were passivated using polyethylene glycol methyl ether thiol (PEG-thiol, average MW 800 Da, 729108, Sigma Aldrich) as previously described^1,3^. The double nanohole samples were immersed in a solution comprised of 2 mM PEG-thiol in ethanol and left overnight (∼18 hours) before being rinsed thoroughly with ethanol and dried using an air gun. Solutions were freshly prepared before each use.

### Nanoaperture optical tweezers setup

Optical components were purchased from Thorlabs as previously described^1–3^. A half-wave plate adjusts the polarisation of the 852 nm laser (Thorlabs, FPL852) to be across the pointed edges of the gap between the two nanoholes (*y-*axis, Fig. 1a)^1,4^. The laser was collimated and expanded to 5 mm in diameter and then focused onto the DNH using a 100× objective (1.25 NA PLN100XO, Olympus). The laser power on the DNH was around 20 mW correlating to around 37^°^C at the focusing point due to laser heating (Figs. S3). Light passing through the sample was collected with a 4× objective (0.1 NA PLN4XP, Olympus), then was focused onto an avalanche photodiode (APD120A/M, Thorlabs), which converted the light intensity to a voltage signal.

### Data acquisition

The avalanche photodiode (APD120A/M, Thorlabs) has a bandwidth of 50 MHz. However, considering the signal bandwidth of the system (∼10 kHz, Fig. 3c and 3d) and to optimise the file size, the voltage signal was recorded at a sampling rate of 1 MHz using a data acquisition card (USB-6361, NI) controlled by a custom LabVIEW program. Based on the Nyquist frequency, this system provides a the theoretical time resolution of 2 μs.

### Microfluidics system

Flow cells were printed using a FormLab 2 printer with Clear V4 resin at a resolution of 50 μm (FormLabs Inc, USA), the same as previously described^1–3^. Two component silicone glue (Twinsil, Picodent, Germany) was used to seal the DNH structure within the flow cell with a 0.17 mm thick glass coverslip. A 50 μm thick piece of double-sided tape (Arcare92712, Adhesive Research, Inc) was used to separate the DNH and glass coverslip, creating a chamber with a volume of 3.5 μL. Flow rate and flow direction were controlled using a syringe pump (Harvard Apparatus, US) through a 12-valve distributor (MUX Distributor, Elveflow, France). For Sam68 N-terminal experiments, the RNA solution is infused to the chamber at a flow rate of ∼1-2 μL/min while Sam68 N-terminal was trapped. Given the ∼10 μL dead volume of the intake tubing, this flow rate required ∼5-10 minutes for RNA solution to arrive at the trapping site.

### Protein and RNA preparation

Human haemoglobin (H7379, Sigma Aldrich), human native tau-441 (T0576, Sigma Aldrich), human GSK3β-tau-441 (SRP0689, Sigma Aldrich), bovine ribonuclease A (R5500, Sigma Aldrich), and bovine serum albumin (A8531, Sigma Aldrich) were prepared in a filtered buffer solution of 0.1 M bis-tris propane, 150 mM NaCl and 20% glycerol at pH 7.2. Bovine actin (A3653, Sigma Aldrich) was prepared in 0.1 M bis-tris propane, 0.2 mM CaCl_2_, 0.2 mM ATP and 20% glycerol at pH 7.2.

Human Sam68 N-terminal (amino acids 1-96) and C-terminal (amino acids 267-368) sequences were cloned and produced at the University of Leicester and comprised as previously described^5^ (sequence also available in Fig. S15a). G8.5 RNA sequence was bought from Dharmacon, Horizon Discovery and is the same as previously described^6^ (sequence also available in Fig. S15b). Both protein solutions were prepared in a filtered solution containing 50 mM sodium phosphate and 150 mM NaCl at pH 6.8.

Proteins were aliquoted into 1 μM aliquots of 100 μL volume and immediately flash frozen in liquid nitrogen before being stored at -20°C, except for GSK3β-tau which was stored at -80^°^C. The proteins were slow thawed on wet ice on the day of use for an experiment.

### Data analysis

Custom MATLAB scripts were used to analyse all the data in this work.

#### Data filtering

Raw data were filtered using a zero-phase Gaussian low-pass filter to the desired cut-off frequency by using the **filtfilt.m** function.

#### Normalisation of optical transmission traces

We used normalised transmission intensity, Δ*I/I*_*0*_, to quantify the relative transmission change upon trapping a single protein. Since the optical signal was recorded by the avalanche photodiode as voltage (V), with Δ*I/I*_*0*_ calculated as Δ*I/I*_*0*_ = (*V – V*_*0*_)/ *V*_*0*_. For trapping traces shown in Fig. 2a-c, *V*_*0*_ is the mean value of the baseline, whilst for the trapped transmission traces (Fig. 2d and Fig. 3a-b), *V*_*0*_ corresponds to the mean value of the trace.

#### Probability density function (PDF)

We filtered the 20-s trace with a cutoff frequency of 10 kHz and calculated the PDF by estimating the kernel density using the **ksdensity.m** function with 300 points. This cutoff frequency was chosen based on the PSD analysis (Fig. 3c), which shows that protein-induced signal variations remained distinguishable from empty DNH noise up to 10 kHz.

#### Trace detrending

To allow well-assigned levels for step fitting, we removed the linear drift of the whole trace using the **detrend.m** function from MATLAB 2022b, as shown in Fig. S16.

#### Deconvolution of PDF and energy landscapes

See details in Supplementary SI-9.

#### Power spectral density (PSD)

To estimate the PSD of the time-domain signal across the trapping trace, we used the sampling frequency (*f*) and the signal vector (*XXX*). The frequency vector was taken over half of the frequency spectrum, from 0 to *f*/2 for the Nyquist frequency using a linearly spaced grid with the number of data points (*N*)/2 size. The power spectrum was then computed using the Fast Fourier Transform (FFT) and the squared magnitude was obtained using |FFT(*XXX*)|^2^ before normalisation by *N* multiplied by *f* as shown in Eq. 1:

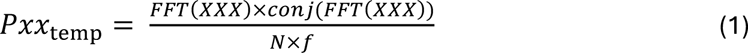

To remove negative frequency components and conserve the total power in the spectrum, the power spectrum was truncated to *N*/2 points and multiplied by 2 as shown in Eq. 2:

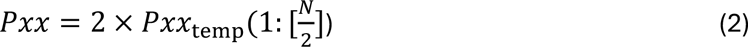

### Finite-difference time-domain simulations

The transmission of DNH structures were modelled based on finite-difference time-domain (FDTD) using a commercial software (Lumerical, Ansys). See Tables S1, S2, and S3 in Supplementary SI-2 for parameters.

#### DNH model structure

See Supplementary SI-2 for details.

#### Volume versus transmission changes

The relationship between particle volume and the change in transmission (Fig. S2e) were simulated by placing a spherical particle (refractive index *n* = 2.0) with radii ranging from 2 to 6 nm in the centre of the DNH (*x* = 0, *y* = 0, *z* = 8).

### Laser heating Simulation

The laser heating simulation is the same as described previously^1^, with details listed in Fig. S3.

## Notes

### Competing Interest Statement

The authors have declared no competing interest.

### Summary of Updates

Altered manuscript title. Formatting changes applied.

## References

1. Holehouse, A. S. & Kragelund, B. B. The molecular basis for cellular function of intrinsically disordered protein regions. Nat Rev Mol Cell Biol 25, 187–211 (2024).

2. Snead, D. & Eliezer, D. Intrinsically disordered proteins in synaptic vesicle trafficking and release. Journal of Biological Chemistry 294, 3325–3342 (2019).

3. Melo, A. M. et al. A functional role for intrinsic disorder in the tau-tubulin complex. Proceedings of the National Academy of Sciences 113, 14336–14341 (2016).

4. Tesei, G. et al. Conformational ensembles of the human intrinsically disordered proteome. Nature 626, 897–904 (2024).

5. Coskuner-Weber, O., Mirzanli, O. & Uversky, V. N. Intrinsically disordered proteins and proteins with intrinsically disordered regions in neurodegenerative diseases. Biophys Rev 14, 679–707 (2022).

6. Colom-Cadena, M. et al. Synaptic oligomeric tau in Alzheimer’s disease — A potential culprit in the spread of tau pathology through the brain. Neuron 111, 2170-2183.e6 (2023).

7. Calabresi, P. et al. Alpha-synuclein in Parkinson’s disease and other synucleinopathies: from overt neurodegeneration back to early synaptic dysfunction. Cell Death Dis 14, 1–16 (2023).

8. Shim, K. H., Kang, M. J., Youn, Y. C., An, S. S. A. & Kim, S. Alpha-synuclein: a pathological factor with Aβ and tau and biomarker in Alzheimer’s disease. Alz Res Therapy 14, 201 (2022).

9. Pan, L., Meng, L., He, M. & Zhang, Z. Tau in the Pathophysiology of Parkinson’s Disease. J Mol Neurosci 71, 2179–2191 (2021).

10. Iakoucheva, L. M., Brown, C. J., Lawson, J. D., Obradović, Z. & Dunker, A. K. Intrinsic Disorder in Cell-signaling and Cancer-associated Proteins. Journal of Molecular Biology 323, 573–584 (2002).

11. Sumithra, B., Saxena, U. & Das, A. B. A comprehensive study on genome-wide coexpression network of KHDRBS1/Sam68 reveals its cancer and patient-specific association. Sci Rep 9, 11083 (2019).

12. Dyson, H. J. & Wright, P. E. Intrinsically unstructured proteins and their functions. Nat Rev Mol Cell Biol 6, 197–208 (2005).

13. Uversky, V.N. Natively unfolded proteins: A point where biology waits for physics. Protein Science 11, 739–756 (2002).

14. Fuxreiter, M. & Tompa, P. Fuzzy Complexes: A More Stochastic View of Protein Function. in Fuzziness: Structural Disorder in Protein Complexes (eds. Fuxreiter, M. & Tompa, P.) 1–14 (Springer US, New York, NY, 2012). doi:10.1007/978-1-4614-0659-4_1.

15. Alderson, T. R., Pritišanac, I., Kolarić, Đ., Moses, A. M. & Forman-Kay, J. D. Systematic identification of conditionally folded intrinsically disordered regions by AlphaFold2. Proceedings of the National Academy of Sciences 120, e2304302120 (2023).

16. Dyson, H. J. & Wright, P. E. NMR illuminates intrinsic disorder. Current Opinion in Structural Biology 70, 44–52 (2021).

17. Bernadó, P. & Svergun, D. I. Structural analysis of intrinsically disordered proteins by small-angle X-ray scattering. Mol. BioSyst. 8, 151–167 (2011).

18. Mukhopadhyay, S. The Dynamism of Intrinsically Disordered Proteins: Binding-Induced Folding, Amyloid Formation, and Phase Separation. J. Phys. Chem. B 124, 11541–11560 (2020).

19. DePristo, M. A., Bakker, P.I.W.de & Blundell, T. L. Heterogeneity and Inaccuracy in Protein Structures Solved by X-Ray Crystallography. Structure 12, 831–838 (2004).

20. Yan, Z. et al. Structure of the rabbit ryanodine receptor RyR1 at near-atomic resolution. Nature 517, 50–55 (2015).

21. Nwanochie, E. & Uversky, V. N. Structure Determination by Single-Particle Cryo-Electron Microscopy: Only the Sky (and Intrinsic Disorder) is the Limit. International Journal of Molecular Sciences 20, 4186 (2019).

22. Punjani, A. & Fleet, D. J. 3DFlex: determining structure and motion of flexible proteins from cryo-EM. Nat Methods 20, 860–870 (2023).

23. Schuler, B., Lipman, E. A. & Eaton, W. A. Probing the free-energy surface for protein folding with single-molecule fluorescence spectroscopy. Nature 419, 743–747 (2002).

24. Schuler, B., Soranno, A., Hofmann, H. & Nettels, D. Single-Molecule FRET Spectroscopy and the Polymer Physics of Unfolded and Intrinsically Disordered Proteins. Annual Review of Biophysics 45, 207–231 (2016).

25. Nettels, D. et al. Single-molecule FRET for probing nanoscale biomolecular dynamics. Nat Rev Phys 6, 587–605 (2024).

26. Ferreon, A. C. M., Gambin, Y., Lemke, E. A. & Deniz, A. A. Interplay of α-synuclein binding and conformational switching probed by single-molecule fluorescence. Proceedings of the National Academy of Sciences 106, 5645–5650 (2009).

27. Neupane, K., Solanki, A., Sosova, I., Belov, M. & Woodside, M. T. Diverse Metastable Structures Formed by Small Oligomers of α-Synuclein Probed by Force Spectroscopy. PLOS ONE 9, e86495 (2014).

28. Berkovich, R. et al. Rate limit of protein elastic response is tether dependent. Proceedings of the National Academy of Sciences 109, 14416–14421 (2012).

29. Sánchez-Rico, C., Voith von Voithenberg, L., Warner, L., Lamb, D. C. & Sattler, M. Effects of Fluorophore Attachment on Protein Conformation and Dynamics Studied by spFRET and NMR Spectroscopy. Chemistry – A European Journal 23, 14267–14277 (2017).

30. Bondos, S. E., Dunker, A. K. & Uversky, V. N. Intrinsically disordered proteins play diverse roles in cell signaling. Cell Commun Signal 20, 20 (2022).

31. Pang, Y. & Gordon, R. Optical Trapping of a Single Protein. Nano Lett. 12, 402–406 (2012).

32. Juan, M. L., Gordon, R., Pang, Y., Eftekhari, F. & Quidant, R. Self-induced back-action optical trapping of dielectric nanoparticles. Nature Phys 5, 915–919 (2009).

33. Letwin, K., Peters, M. & Gordon, R. Conformational Stability at Low Temperatures Using Single Protein Nanoaperture Optical Tweezers. J. Phys. Chem. B 129, 2402–2407 (2025).

34. Jiang, Q., Rogez, B., Claude, J.-B., Baffou, G. & Wenger, J. Temperature Measurement in Plasmonic Nanoapertures Used for Optical Trapping. ACS Photonics 6, 1763–1773 (2019).

35. Yang-Schulz, A. et al. Direct observation of small molecule activator binding to single PR65 protein. npj Biosensing 2, 1–10 (2025).

36. Kotnala, A. & Gordon, R. Double nanohole optical tweezers visualize protein p53 suppressing unzipping of single DNA-hairpins. Biomed. Opt. Express, BOE 5, 1886–1894 (2014).

37. Ying, C. et al. Watching Single Unmodified Enzymes at Work. Preprint at 10.48550/arXiv.2107.06407 (2021).

38. Yousefi, A. et al. Optical Monitoring of In Situ Iron Loading into Single, Native Ferritin Proteins. Nano Letters 23, 3251–3258 (2023).

39. Yousefi, A. et al. Structural Flexibility and Disassembly Kinetics of Single Ferritin Molecules Using Optical Nanotweezers. ACS Nano 18, 15617–15626 (2024).

40. Peters, M. et al. Energy landscape of conformational changes for a single unmodified protein. npj Biosensing 1, 1–10 (2024).

41. Booth, L. S. et al. Modelling of the dynamic polarizability of macromolecules for single-molecule optical biosensing. Sci Rep 12, 1995 (2022).

42. Quinten Michael. Beyond Mie’s Theory I – Nonspherical Particles. in Optical Properties of Nanoparticle Systems 255 (John Wiley & Sons, Ltd, 2011). doi:10.1002/9783527633135.ch9.

43. Zargarbashi, S., Xu, L., Mellor, C. J., Rahmani, M. & Ying, C. Monitoring Conformational Dynamics of Single Unmodified Proteins using Plasmonic Nanotweezers. Journal of Visualized Experiments (JoVE) e68093 (2025) doi:10.3791/68093.

44. Li, L., Li, C., Zhang, Z. & Alexov, E. On the Dielectric “Constant” of Proteins: Smooth Dielectric Function for Macromolecular Modeling and Its Implementation in DelPhi. J. Chem. Theory Comput. 9, 2126–2136 (2013).

45. Sarimov, R. M., Matveyeva, T. A. & Binhi, V. N. Laser interferometry of the hydrolytic changes in protein solutions: the refractive index and hydration shells. J Biol Phys 44, 345–360 (2018).

46. Schuler, B. & Hofmann, H. Single-molecule spectroscopy of protein folding dynamics— expanding scope and timescales. Current Opinion in Structural Biology 23, 36–47 (2013).

47. Abramson, J. et al. Accurate structure prediction of biomolecular interactions with AlphaFold 3. Nature 630, 493–500 (2024).

48. Meng, E. C. et al. UCSF ChimeraX: Tools for structure building and analysis. Protein Science 32, e4792 (2023).

49. Maslov, I. et al. Sub-millisecond conformational dynamics of the A2A adenosine receptor revealed by single-molecule FRET. Commun Biol 6, 1–15 (2023).

50. Woodside, M. T. & Block, S. M. Reconstructing Folding Energy Landscapes by Single-Molecule Force Spectroscopy. Annu Rev Biophys 43, 19–39 (2014).

51. Chong, S.-H. & Ham, S. Folding Free Energy Landscape of Ordered and Intrinsically Disordered Proteins. Sci Rep 9, 14927 (2019).

52. Strodel, B. Energy Landscapes of Protein Aggregation and Conformation Switching in Intrinsically Disordered Proteins. Journal of Molecular Biology 433, 167182 (2021).

53. Viegas, R. G., Martins, I. B. S. & Leite, V. B. P. Understanding the Energy Landscape of Intrinsically Disordered Protein Ensembles. J. Chem. Inf. Model. 64, 4149–4157 (2024).

54. Burger, V. M., Gurry, T. & Stultz, C. M. Intrinsically Disordered Proteins: Where Computation Meets Experiment. Polymers 6, 2684–2719 (2014).

55. Marsh, J. A. & Forman-Kay, J. D. Sequence Determinants of Compaction in Intrinsically Disordered Proteins. Biophysical Journal 98, 2383–2390 (2010).

56. Jin, F. & Gräter, F. How multisite phosphorylation impacts the conformations of intrinsically disordered proteins. PLOS Computational Biology 17, e1008939 (2021).

57. Adzhubei, A. A., Sternberg, M. J. E. & Makarov, A. A. Polyproline-II Helix in Proteins: Structure and Function. Journal of Molecular Biology 425, 2100–2132 (2013).

58. Chebrek, R., Leonard, S., de Brevern, A. G. & Gelly, J.-C. PolyprOnline: polyproline helix II and secondary structure assignment database. Database 2014, bau102 (2014).

59. Lukong, K. E. & Richard, S. Sam68, the KH domain-containing superSTAR. Biochimica et Biophysica Acta (BBA) - Reviews on Cancer 1653, 73–86 (2003).

60. Lin, Q., Taylor, S. J. & Shalloway, D. Specificity and Determinants of Sam68 RNA Binding: IMPLICATIONS FOR THE BIOLOGICAL FUNCTION OF K HOMOLOGY DOMAINS *. Journal of Biological Chemistry 272, 27274–27280 (1997).

61. Feracci, M. et al. Structural basis of RNA recognition and dimerization by the STAR proteins T-STAR and Sam68. Nat Commun 7, 10355 (2016).

62. Malki, I. et al. Cdk1-mediated threonine phosphorylation of Sam68 modulates its RNA binding, alternative splicing activity and cellular functions. Nucleic Acids Research 50, 13045– 13062 (2022).

63. Ozdilek, B. A. et al. Intrinsically disordered RGG/RG domains mediate degenerate specificity in RNA binding. Nucleic Acids Research 45, 7984–7996 (2017).

## Materials and methods reference list

1. Ying, C. et al. Watching Single Unmodified Enzymes at Work. Preprint at 10.48550/arXiv.2107.06407 (2021).

2. Yousefi, A. et al. Optical Monitoring of In Situ Iron Loading into Single, Native Ferritin Proteins. Nano Lett. 23, 3251–3258 (2023).

3. Zargarbashi, S., Xu, L., Mellor, C. J., Rahmani, M. & Ying, C. Monitoring Conformational Dynamics of Single Unmodified Proteins using Plasmonic Nanotweezers. Journal of Visualized Experiments (JoVE) e68093.(2025) doi:10.3791/68093.

4. Ghorbanzadeh, M., Jones, S., Moravvej-Farshi, M. K. & Gordon, R. Improvement of Sensing and Trapping Efficiency of Double Nanohole Apertures via Enhancing the Wedge Plasmon Polariton Modes with Tapered Cusps. ACS Photonics 4, 1108–1113 (2017).

5. Malki, I. et al. Cdk1-mediated threonine phosphorylation of Sam68 modulates its RNA binding, alternative splicing activity and cellular functions. Nucleic Acids Research 50, 13045–13062 (2022).

6. Lin, Q., Taylor, S. J. & Shalloway, D. Specificity and Determinants of Sam68 RNA Binding: IMPLICATIONS FOR THE BIOLOGICAL FUNCTION OF K HOMOLOGY DOMAINS *. Journal of Biological Chemistry 272, 27274–27280 (1997).

