## Supplementary Information for "Direct observation of conformational dynamics in intrinsically disordered proteins at the single-molecule level"

### SI-1 Nanoaperture optical tweezers setup

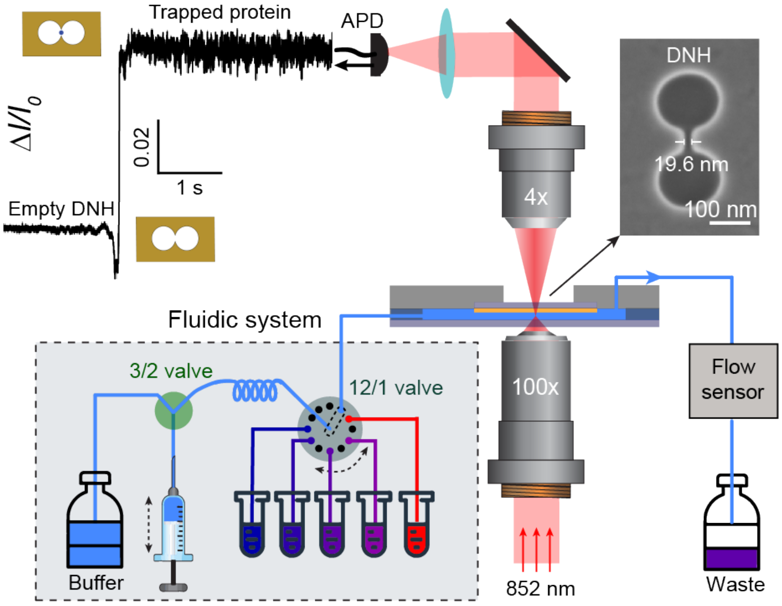


**Fig. S1|** Schematic of the nanoaperture optical tweezers setup. A Collimated laser at 852 nm is focused onto the double nanohole (DNH, with SEM image on the right) structure by a 100x objective (NA 1.25) to excite localised surface plasmonic resonance (LSPR). Laser transmitted through the DNH is subsequently collected by a 4x objective (NA 0.1) and detected by an avalanche photodiode (APD) to record the transmission intensity over time. **Top left inset:** transmission intensity trace change upon trapping a protein. Increased fluctuations in the normalised transmission intensity ($\Delta I/I_0$) signify the trapping of a protein. **Bottom left inset:** incorporated microfluidics system allows altering solution conditions whilst the protein is trapped. See materials and methods for details.

#### SI-2 Finite-difference time-domain (FDTD) simulations

To understand the relationship between transmission intensity changes, $\Delta I/I_0$, and the shape/volume of the trapped particle, we conducted finite-difference time-domain (FDTD) simulations using commercial software (Lumerical, Ansys). **Figures. S2a–c** confirms the strong field enhancement localised in the DNH gap due to localised surface plasmon resonance (LSPR) excitation. Introduction of a spherical particle (edge positioned ~2.5 nm from the gold surface, **Fig. S2c**) further enhances the local field intensity, thereby stabilising the trapping of the particle.

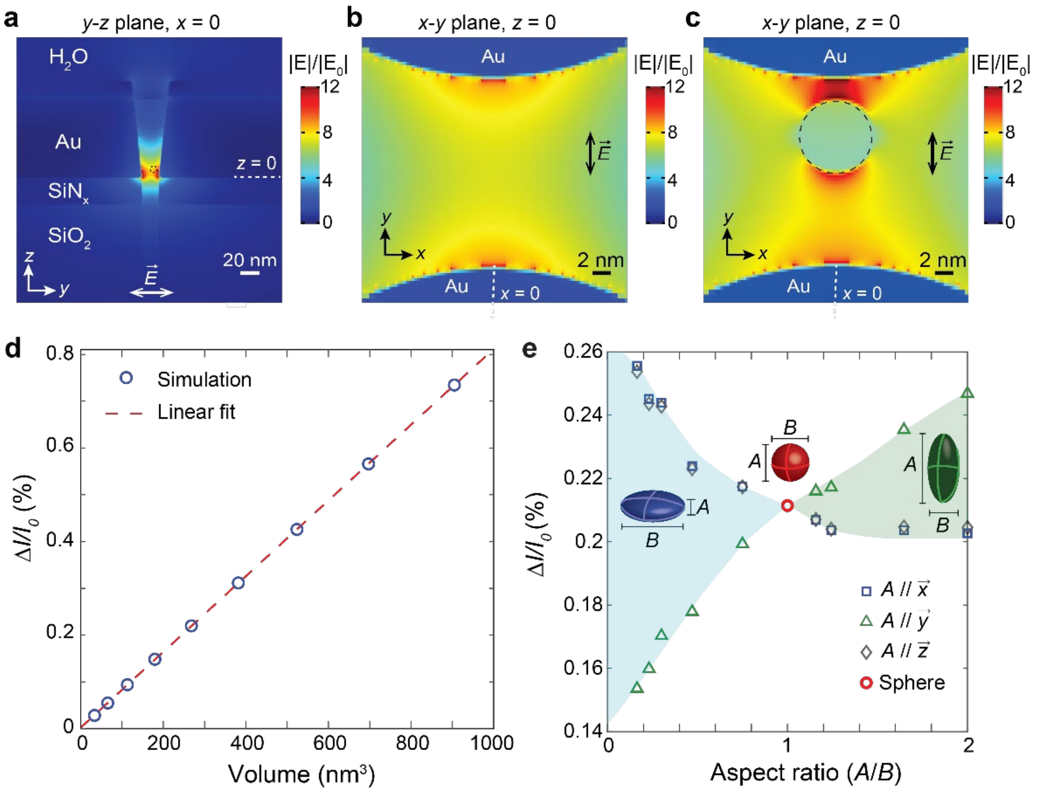


**Fig. S2| a**, Field distribution in an empty DNH on the *y-z* plane at $x = 0$. **b**,**c**, Zoomed field enhancement of an empty DNH (**b**) and a DNH containing a spherical particle (**c**) (refractive index, $n = 2$, radius, $r = 4$ nm) at $z = 0$. For each simulation the incident laser (852 nm ± 10 nm) is polarised along the *y*-axis. **d,** Linear relationship between the change in normalised transmission intensity, $\Delta I/I_0$, and volume of spherical particles ($n = 2$) within the DNH gap, as indicated by the dashed circle in **a**, along with the linear fit (red dashed line). Spheres with 9 different radii, positioned with an edge-to-edge distance of 2.5 nm from the DNH, were simulated with parameters detailed in Table S1. **e**, Orientation- and shape-dependency of $\Delta I/I_0$ for ellipsoids with various aspect ratios (*A*/*B*), where ellipsoid shape axes are defined as (*A, B, B*), with $A/B < 1$ corresponding to oblate, $A/B = 1$ to sphere, and $A/B > 1$ to prolate (see Supplementary Table S2 for detailed parameters). Three orientations were considered: *A*-axis along the *x*- (blue squares), *y*- (green triangles) and *z*-axes (grey circles).

**Figures S2d and S2e** present the simulated effects of particle volume, shape and orientation on $\Delta I/I_0$. The presence of the nanoparticle alters the refractive index within the DNH gap, leading to changes in transmission intensity, $\Delta I/I_0$. The refractive index of a nanoparticle depends on its polarisability, which is associated with volume, shape anisotropy, and dielectric constant[1]. The FDTD simulations reveal a linear relationship between $\Delta I/I_0$ and spherical particle volume (**Fig. S2d,** the particle parameters used are listed in **Table S1**). This relationship suggests that a larger protein produces a greater $\Delta I/I_0$ signal when trapped in the same DNH structure. The influence of ellipsoidal geometry on $\Delta I/I_0$ is complex, as it depends on both the aspect ratio and the orientation relative to the DNH. **Fig. S2e** demonstrates how $\Delta I/I_0$ changes with aspect ratio and orientation of the ellipsoid (see **Table S2** for particle parameters). For spherical particles (aspect ratio = 1), no orientation-dependent signal changes occur, therefore, we expect stable $\Delta I/I_0$ with non-system fluctuations primarily arising from translational diffusion within the trapping well. For non-spherical geometries (aspect ratio ≠ 1), depending on the orientation, $\Delta I/I_0$ can either increase or decrease relative to an equivalent volume sphere. While rotational diffusion would theoretically cause larger $\Delta I/I_0$ fluctuations for non-spherical particles, we consistently observe greater, yet stable, transmission changes for globular proteins in extended conformations compared to compact states[2–4]. The absence of increased fluctuations in $\Delta I/I_0$ might be attributed to the optimal alignment of ellipsoidal particles, as the self-induced back action trapping mechanism preferentially stabilises orientations that maximise $\Delta I/I_0$.

**Table S1.** Particle parameters used for Lumerical simulation in Fig. S2d.

| Radius (nm) | Volume (nm$^3$) |
|---|---|
| 2.0 | 33.51 |
| 2.5 | 65.45 |
| 3.0 | 113.10 |
| 3.5 | 179.59 |
| 4.0 | 268.08 |
| 4.5 | 381.70 |
| 5.0 | 523.60 |
| 5.5 | 696.91 |
| 6.0 | 904.78 |

**Table S2**. Parameters for ellipsoid dimensions for Lumerical simulation in Fig. S2e.

| *A* (nm) | *B* (nm) | Aspect ratio (*A*/*B*) | Volume (nm$^3$) |
|---|---|---|---|
| 2.4 | 14.6 | 0.164 | 267.86 |
| 3.0 | 13.0 | 0.231 | 265.46 |
| 3.6 | 12.0 | 0.3 | 271.43 |
| 4.8 | 10.2 | 0.471 | 261.48 |
| 6.6 | 8.8 | 0.75 | 267.61 |
| 8.0 | 8.0 | 1.0 | 268.08 |
| 8.8 | 7.6 | 1.158 | 266.14 |
| 9.2 | 7.4 | 1.243 | 263.78 |
| 11.2 | 6.8 | 1.64 | 271.10 |
| 12.8 | 6.4 | 2.0 | 274.52 |

The DNH was illuminated with a Gaussian shaped plane-wave source (90° polarisation, *z*-axis in the backward direction, 1 µm spot size, $\lambda$ = 852 nm ± 10 nm, relative $z$ = 80 nm). Three mesh sizes were used for the simulation (see **Table S3** for details). Simulations used perfectly matched layer (PML) boundary conditions with a simulation time of 5000 fs and temperature of 310.1 K. Trapping experiments used the laser power corresponding to a local temperature of 37°C (**Figs. S3a and S3b**). The DNH structure substrate is comprised of a 100 nm Au layer (Johnson and Christy), 5 nm Ti (Palik), and 30 nm $SiN_x$ (refractive index, $n$ = 2) on a $SiO_2$

substrate layer (Palik). The DNH geometry was formed using an SEM image as previously described[3]. The tapering of the DNH, observed in the SEM images in **Fig. S4**, was accounted for by using truncated m-sided pyramids (*m* = 6), with *x*-span = 50 nm, *y*-span = 130 nm, *z*-span = 100 nm (200 slices per pyramid). The coordinate origin (*x*, *y*, *z*) = (0, 0, 0) was defined at the centre of the Au/Ti interface at the bottom of the DNH gap. Transmission values were collected using a frequency domain monitor at *z* = 8 nm with a 24 × 24 $nm^2$ *xy*-plane detection area. The overall and zoomed resonance in the DNH was imaged with frequency domain monitors (overall: 2D *x*-normal, *x* = 0 nm, *y*-span = 300 nm, *z*-span = 300 nm, zoomed region: 2D *z*-normal, *x* = 0 nm, *x*-span = 30 nm, *y* = 0 nm, *y*-span = 30 nm and *z* = 8 nm).

**Table S3.** Lumerical simulation mesh parameters for Supplementary Fig. S2.

| Mesh | Parameter | Value | Details |
|---|---|---|---|
| FDTD region | *Mesh type* | Auto non-uniform | Generates general mesh based on mesh accuracy |
| | *Conformational mesh* | Conformational variant 0 | Refines mesh to improve accuracy |
| | *Mesh accuracy* | 5 | Accuracy of mesh |
| | *Background* $n$ | 1.35 | Default refractive index of objects in simulation |
| | *Relative xy* | 0 nm | *xy* position relative to origin |
| | *Relative z* | -500 nm | *z* position relative to origin |
| | *xy-span* | 1400 nm | Range of *xy*-axes from *relative xy* |
| | *z-span* | 1500 nm | Range of *z*-axis from *relative* z |
| | *Min mesh step* | 0.1 nm | Minimum mesh size |
| Substrate mesh | *Max xyz mesh step* | 10 nm | Maximum mesh size for Yee cells in *xyz* dimensions |
| | *Relative xy* | 0 nm | *x* and *y* position relative to origin |
| | *Relative z* | 65 nm | *z* position relative to origin |
| | *x-span* | 600 nm | Range of *x*-axis from *relative x* |
| | *y-span* | 500 nm | Range of *y*-axis from *relative y* |
| | *z-span* | 370 nm | Range of *z*-axis from *relative z* |
| DNH mesh | *Max xyz mesh step* | 0.5 nm[†]<br>0.3 nm[††] | Maximum mesh size for Yee cells in *xyz* dimensions |
| | *Relative xy* | 0 nm | *xy* position relative to origin |
| | *Relative z* | 12 nm | *z* position relative to origin |
| | *xy-span* | 24 nm | Range of *xy*-axes from *relative xy* |
| | *z-span* | 30 nm | Range of *z*-axis from *relative* z |

$n$, refractive index.

[†]Parameter for volume vs transmission simulation (Fig. S2d).

[††]Parameter for shape and orientation simulation (Fig. S2e).

### SI-3 Extended protein conformations increase the local refractive index

The refractive index ($n$) can be described by the Lorentz-Lorenz equation[5] for a solution containing $N$ particles, each with a polarisability of $\alpha$, the relationship is given by:

$$\frac{n^2-1}{n^2+2} = \frac{4\pi}{3} N\alpha \quad \text{(S1)}$$

Within the DNH sensing volume (field enhanced zone), containing both solution and protein molecules, **Eq. S1** becomes:

$$\frac{n^2-1}{n^2+2} = \frac{4\pi}{3} (N_p\alpha_p + N_w\alpha_w + N'_w\alpha'_w) \quad \text{(S2)}$$

Where, $N_p$ and $\alpha_p$ are the number and polarisability of amino acids within the protein; $N_w$ and $\alpha_w$ are the number and polarisability of water molecules in the bulk volume; and $N'_w$ and $\alpha'_w$ are the number and polarisability of water molecules in protein cavities or the near-surface layer, comprising the hydration shell of the protein.

The hydration shell exhibits a higher average water density than bulk water due to the electrostatic fields generated by protein atoms, which induce a preferential alignment of water dipoles. When protein molecules unfold—from tightly compact to loosely extended conformations—the number of water molecules in the hydration shell ($N'_w$) increases as water molecules penetrate newly accessible regions[6,7]. Meanwhile, the number of bulk water molecules ($N_w$) and the intrinsic properties of the amino acids remain unchanged. This increase in water density enhances the local refractive index within the sensing volume considering the relationship shown in **Eq. S2**.

For globular proteins, extended conformations yield higher $\Delta I/I_0$ signals compared to their folded states[4,8]. This effect arises from both local refractive index changes (mentioned above) and shape anisotropy at a specific orientation, where the self-induced back-action (SIBA) trapping mechanism induces bias in the protein towards the orientation that maximises transmission intensity changes. For IDPs, which continuously sample compact and extended conformations on submicrosecond-to-second timescales[9], two additional factors may contribute to the large fluctuations in $\Delta I/I_0$. First, rapid changes in local refractive index due to hydration dynamics lead to significant signal variability. Second, the short residence time for each conformational state may not allow adequate time for electric field-induced orientation bias. Consequently, the trapped IDP randomly samples orientations and conformations, yielding the orientation-dependent $\Delta I/I_0$ variations as illustrated in **Fig. S2e**.

### SI-4 Laser-induced heating

**Figure S3** shows the simulated laser heating profile of the DNH structure obtained through finite element simulations. We performed the simulation using the transient heat transfer model of COMSOL Multiphysics as described in previous work[10]. The laser was focused onto the gold layer with a spot size of 1.36 μm (measured). The absorption coefficient of water and gold at 852 nm were set to be 4.4372 $m^{-1}$ and $7.87 \times 10^{7}$ $m^{-1}$, respectively.

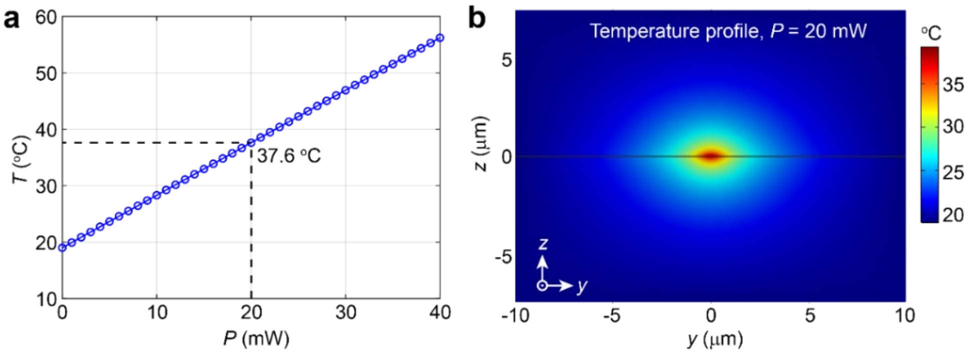


**Fig. S3|** Finite element simulation (FES) of laser heating. Simulated laser heating of the DNH when illuminated by a focused laser beam (852 nm) with a spot size of 1.36 μm (100×, NA 1.25). **a**, Linear relationship between laser power and temperature at the hotspot, with 20 mW marked by dashed line. **b**, Temperature profile of the laser induced heating effect at a laser power of 20 mW. The room temperature was set to 19 °C.

### SI-5 Scanning electron microscopy (SEM) images of representative DNH structures

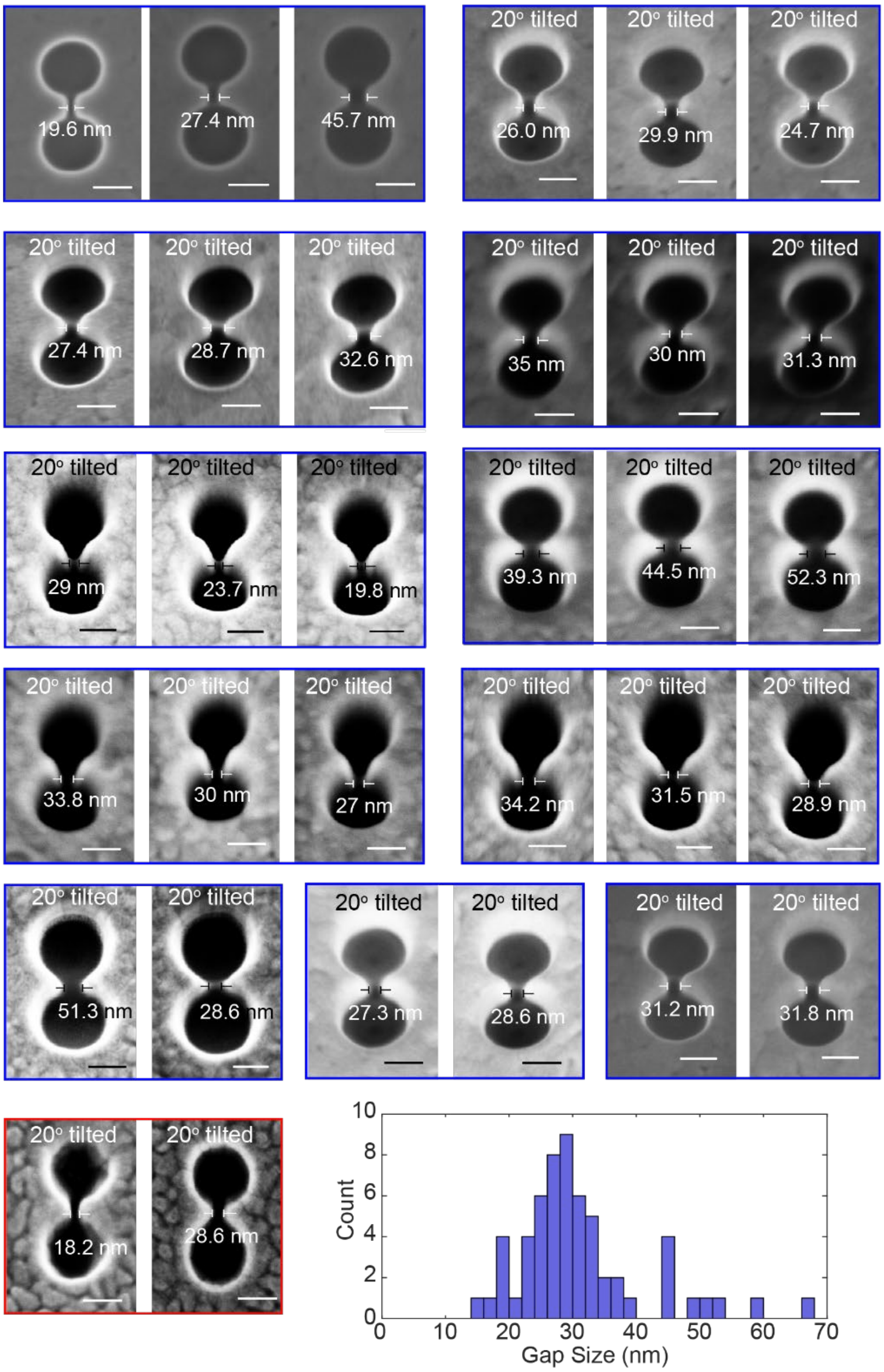


**Fig. S4|** Representative SEM images demonstrating variations in DNH geometries using the same fabrication parameters. Structures from the same chips (blue / red boxes) show smaller gap size deviations compared to gap size differences across chips. DNHs fabricated by a third party (red box) were employed to confirm experimental reproducibility. **Bottom right:** gap size distribution histogram revealing gap sizes predominantly lie within 25-35 nm ($n$ = 61 structures). Note: SEM imaging induces irreversible damage, making the imaged structures no longer suitable for trapping experiments. Therefore, all trapping experiments were conducted using non-imaged structures, assuming comparable geometries as DNHs from the same fabrication batch.

### SI-6 Other trapping traces of IDPs/IDRs and globular proteins of similar molecular weight

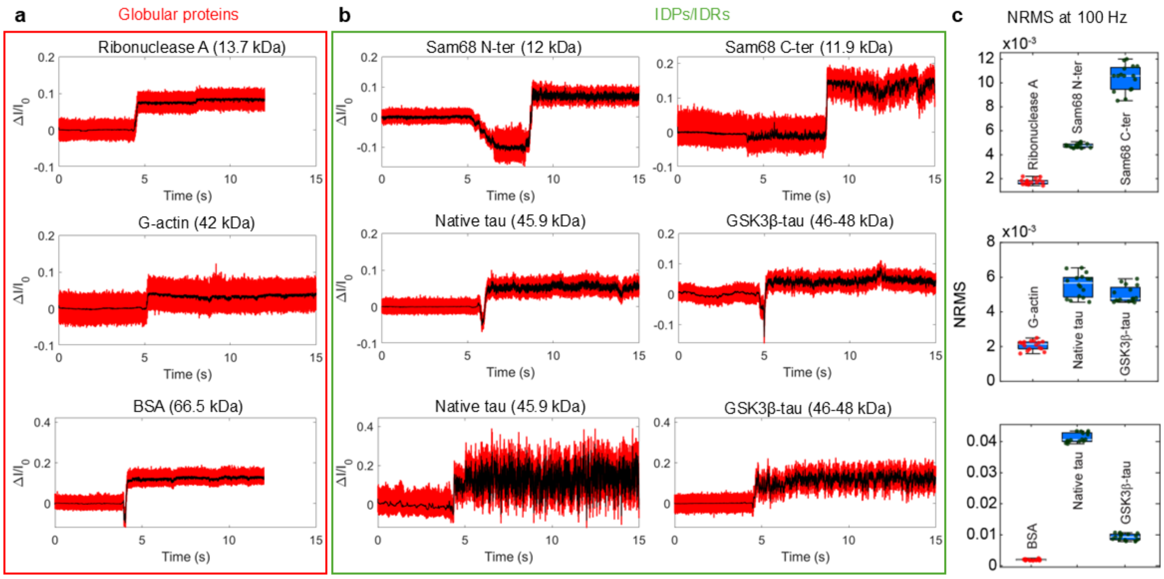


**Fig. S5|** Comparison of trapping traces for globular proteins (**a**, red box) and IDP/IDRs (**b**, green box), along with **c**, a boxplot of normalised root mean square (NRMS) values derived from transmission intensity traces. Trapping traces shown are raw (red) and digitally filtered at 1 kHz (black). The NRMS was calculated from traces filtered at 100 Hz to assess low-frequency fluctuations, using a 5-second sliding window shifted in 1-second increments. At similar molecular weights, IDP/IDRs exhibit greater signal fluctuations during trapping, reflecting their increased structural flexibility compared to globular proteins. Traces were acquired using different double-nanohole (DNH) structures; only those with similar $\Delta I/I_0$ values upon trapping are compared here.

### SI-7 Representative trapping trace showing protein adhesion within the DNH

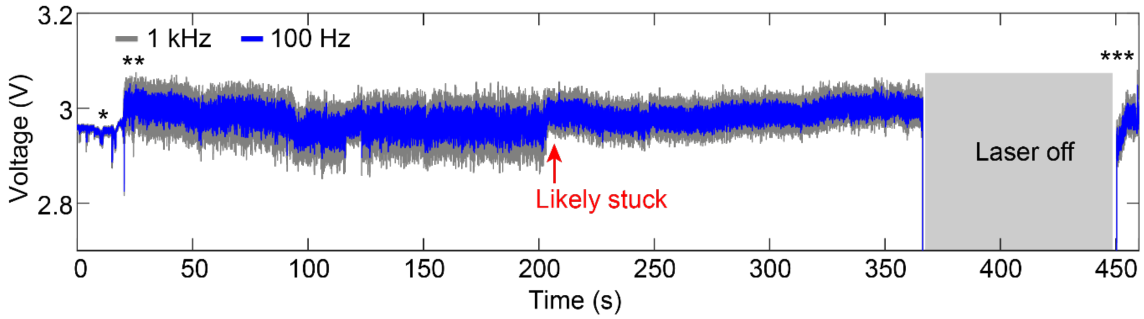


**Fig. S6|** Example trapping trace of GSK3β-tau showing protein adhesion in the DNH gap and does not release after the laser is turned off. Trace was recorded at 1 MHz and is shown filtered to 1 kHz (grey) and 100 Hz (blue). We hypothesise that the protein was stuck at the time indicated by the red arrow, as the reduction in signal fluctuation suggests restricted protein motion, likely due to adhesion to the gold surface. Asterisks indicate the following events: (*): baseline before trapping, (**): trapping event, (***): protein still stuck after turning the laser off.

### SI-8 Raw trapping traces of haemoglobin, GSK3β-tau and native tau-441

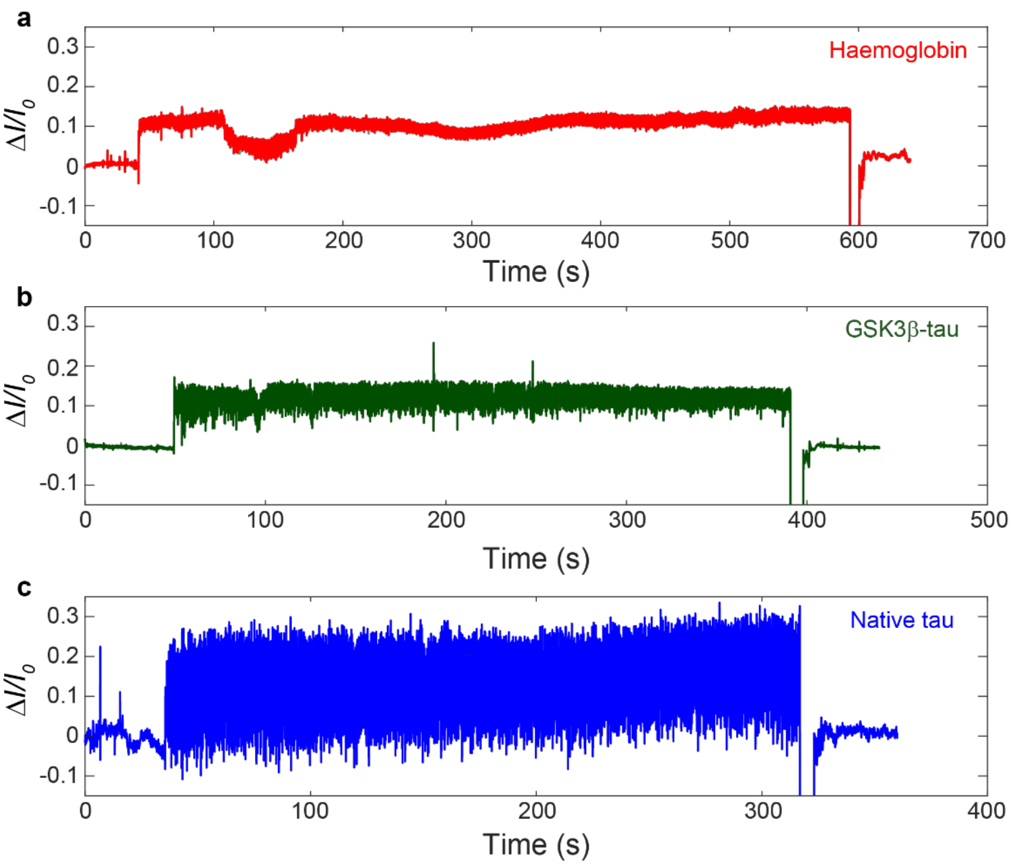


**Fig. S7|** Raw trapping traces of data shown in Fig. 2 for **a**, haemoglobin, **b**, GSK3β-tau and **c**, native tau-441. Data is shown filtered to 1 kHz.

### SI-9 Deconvolution of PDFs and calculation of energy landscapes

All probability density functions (PDFs) in the main text, shaded area of **Figs. 2e, 3c, and 3d**, were calculated from 10 kHz filtered 20-second traces. Within the 10 kHz bandwidth, protein-induced signal variations are distinguishable from empty DNH, as suggested by the PSD in **Fig. 3c**. We deconvoluted these PDFs using a Gaussian point spread function (PSF) via the iterative Lucy-Richardson algorithm (deconvlucy.m), with the PSF as root mean standard deviation (RMSD) of 1 second of haemoglobin trapping.

We obtained the 2D energy landscapes by taking the negative logarithm of the deconvoluted PDF, as described previously[4].

$$E\left(\frac{\Delta I}{I_0}\right) = -k_B T \cdot ln\left[PDF\left(\frac{\Delta I}{I_0}\right)\right] \qquad \text{(S3)}$$

where $k_B$ is the Boltzmann constant, $T$ is the temperature, and $\frac{\Delta I}{I_0}$ is the normalised transmission change.

To visualise multiple conformations, we converted 2D energy landscapes to 3D. To do so, we first fitted the deconvoluted PDF with a multi-peak Gaussian function,

$$G(x) = \sum_{i=1}^{n} A_i e^{-2(\frac{x-\mu_i}{\sigma_i})} \tag{S4}$$

where $A_i$, $\mu_i$ and $\sigma_i$ are the amplitude, peak location and standard deviation of the *i*-th Gaussian peak, respectively. The dashed curve in **Figure S8a** illustrates the multi-peak Gaussian fit to the deconvoluted PDF of GSK3β-tau (also presented in **Figure 2e**), with each peak displayed in different colours. Here, the PDF was intentionally overfitted to maximise accuracy in reconstructing the overall profile, as the fitting parameters are not intended to carry physical meaning.

We then converted the fitted Gaussian function to the energy landscape using

$$U(x) = -k_B T \cdot \ln\,[G(x)] \tag{S5}$$

where, the troughs in the energy landscapes correspond to the peaks in the PDFs. **Figure S8b** shows that the fitted energy landscape (black dashed curve) closely matches the measured curve (green curve).

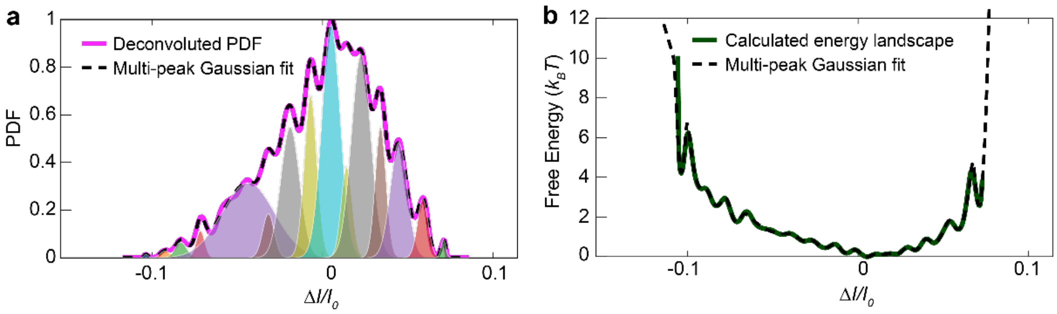


**Fig. S8|** Multi-peak Gaussian fit to **a,** PDF and **b**, conversion to 2D energy landscape. Colour-shaded regions in **a** represent individual fitted peaks. Values for Δ*I*/*I*$_0$ were taken over 20 s and digitally filtered to 10 kHz.

For 3D conversion, we mapped the peaks to a polar axis with radius *R* defined as half the distance between first and last peak locations, $R = \frac{(\mu_N - \mu_1)}{2}$. The angular coordinate $\theta_i$ for the *i*-th peak was distributed between [0.1π, 0.9 π] based on peak position:

$$\theta_i = 0.9\pi\left(1 - \frac{\mu_i - \mu_1}{\mu_N - \mu_1}\right) + 0.1\pi \tag{S6}$$

This angular distribution ensures that different conformations are represented according to their relative position in 2D. The radial coordinate $r_i$ is scaled by the peak amplitude:

$$r_i = \frac{\max(A_i) - A_i}{\max(A_i) - \min(A_i)} R \tag{S7}$$

**Eq. S7** ensures predominant peaks are located near the centre of the 3D plot. **Figure S9** shows peak locations on a polar plot.

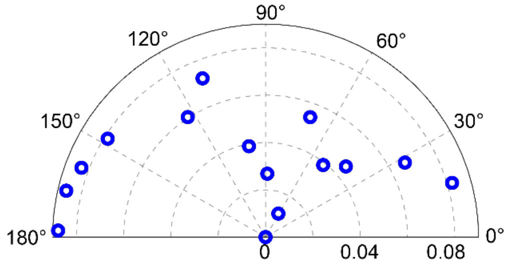


**Fig. S9|** Polar plot of peak locations, with each blue circle representing a fitted peak.

With the peaks being mapped onto a 2D surface, we converted the fitted PDF to a 3D multi-peak Gaussian function, using the peak locations $(\mu_{xi}, \mu_{yi}) = (r_i, \theta_i)$, amplitudes $A_i$ and the standard deviations $\sigma_i$. Finally, the 3D energy landscape (**Fig. 2f and Figs. S10 and S11**) was obtained by taking the negative logarithm of the 3D PDF, with red and blue circles marking fitted troughs.

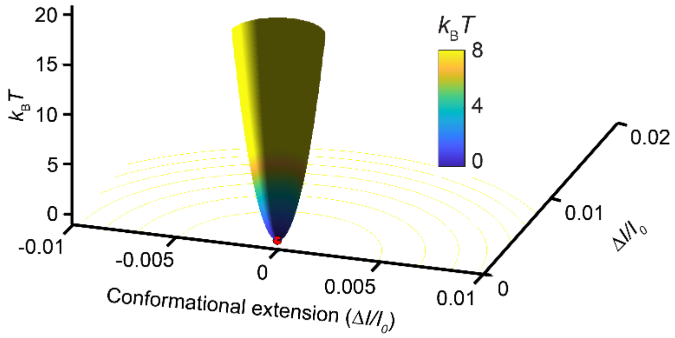


**Fig. S10|** 3D energy landscape of haemoglobin plotted using the deconvoluted PDF in Figure 2e. Funnel shape with a large energy barrier is representative of a typical globular protein with one dominant stable state. A small conformational extension range indicates little change in global shape.

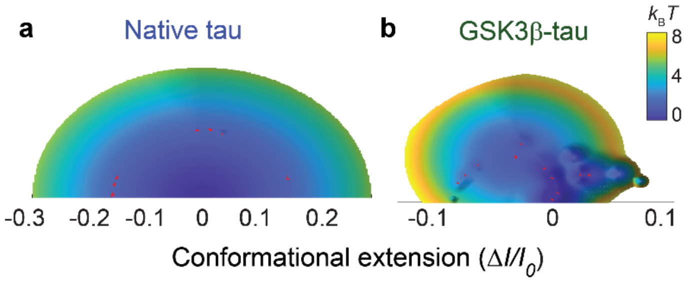


**Fig. S11|** Top-down view of 3D landscapes shown in Fig. 2f for **a**, native tau-441 and **b**, GSK3β-tau. GSK3β-tau displays more troughs than native tau-441, indicating new conformational states not sampled by the unphosphorylated variant.

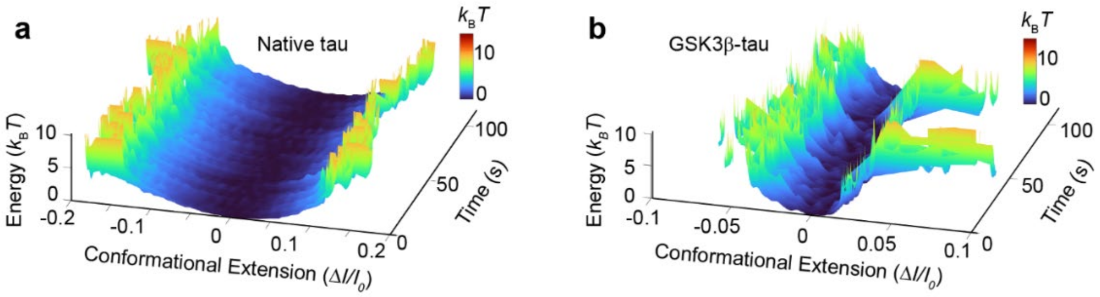


**Fig. S12|** Continuous 3D energy landscapes of IDPs over 120 s. **a**, Native tau-441 and **b**, GSK3β-tau. The energy landscapes were calculated from 1 kHz filtered data using a 5-second window sliding with 100-ms step increments.

**SI-10 Effect of GSK3β induced phosphorylation on tau-441 conformational dynamics**

Post-translational modifications, such as phosphorylation, are of particular interest due to their role in regulating protein function and stability, as well as their contribution to pathological conformational changes in proteins when dysregulated. In tau, phosphorylation regulates its physiological role in microtubule binding *in vivo*[11], while hyperphosphorylation is known to promote aggregation and pathogenic conformations associated with Alzheimer's disease[12].

Native tau-441 contains around 85 putative phosphorylation sites across serine (Ser), threonine (Thr), and tyrosine residues[13]. GSK3β is capable of phosphorylating up to 42 Ser/Thr residues, with site specificity dependent on the presence of a pre-phosphorylated (primed) motif. GSK3β preferentially targets sequences containing Ser/Thr-X-X-X-pSer/pThr and is classified as a proline-directed kinase[14,15], although GSK3β can also phosphorylate unprimed sites[14].

The structural effects of phosphorylation on IDPs depend on both the number and location of modified residues[16], influencing overall disorder, compaction, and the conformational ensemble. The global conformation of an IDP is influenced by its proline content and net charge, with compaction typically being observed as either decreases[17]. Native tau-441 has a theoretical isoelectric point (pI) of 8.24 (**Fig. S13**), and so will present a net positive charge at physiological pH values. As such, by introducing negatively charged phosphate groups via phosphorylation, electrostatic interactions can drive compaction within native tau-441[18].

Previous studies demonstrate that the conformational response of tau-441 to phosphorylation is highly context-dependent. For example, phosphorylation at S202 and T205[19], or pseudo-phosphorylation at S199E, S202E, and T205E or at S396E and S404E[20], has been associated with reduced compaction. Conversely, the combination of these pseudo-phosphorylation sites (S199E, S202E, T205E, S396E, S404E) induced global compaction, which was further increased upon the addition of T212E and S214E[20]. It is important to note that phosphorylated Ser and Thr residues typically present a double charge at physiological pH, than glutamic or aspartic acid (potential -2 vs -1)[21] altering their effects on tau structure.

Our experimental data suggests that phosphorylation of native tau-441 by GSK3β leads to increased conformational order and compaction, as observed in the more defined and stable states sampled by GSK3β-tau compared to native tau-441 (**Fig. 2, Fig. S11**). Two primary mechanisms may underlie this behaviour. First, phosphorylation reduces the net charge of the positively charged tau, promoting compaction. Likely phosphorylation sites for isolated GSK3β include T175, T181, T205, T231, located in the proline rich domain, and S396, S400, and S404, in the C-terminal[22–24] (**Fig. S13**). Although the exact phosphorylation pattern in our commercially obtained GSK3β–tau was not provided, literature suggests that T231 is a key GSK3β phosphorylation site and may be essential for subsequent phosphorylation events, particularly at the C-terminal following local conformational rearrangement at the N-terminal[25]. Second, phosphorylation may induce secondary structure transitions. Residues within the proline-rich domain (I151-Q244) are prone to adopt transient polyproline type II (PPII) helices[26]. Phosphorylation at T231 has been shown to promote α-helical structure formation[27], which are more compact than PPII helices (5.4 Å/turn vs 9.3 Å/turn)[28]. The presence of a stabilising salt bridge between phosphorylated T231 and R230 has also been reported, further increasing conformational order[27]. Similarly, phosphorylation at S396 and

S404 may permit salt bridge formation with local residues such as K395 or R406, leading to increased structural stability.

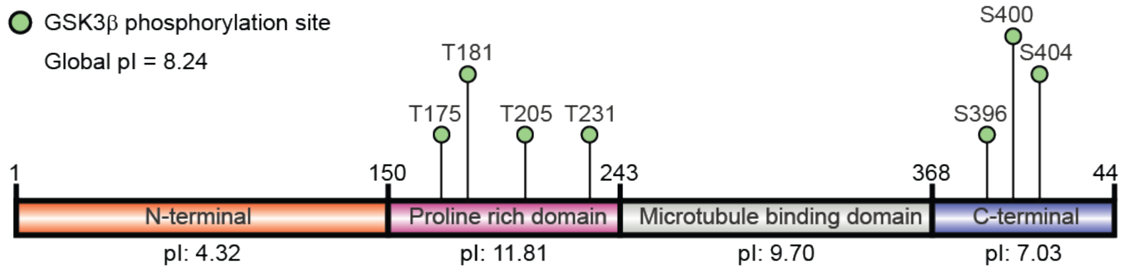


1 MAEPRQEFEVMEDHAGTYGLGDRKDQGGYTMHQDQEGDTDAGLKESPLQTPTEDGSEEPGSETSDAKSTPTAEDVTAPLVDEGAPGKQAAAQPHTEIPEG 100

101 TTAEEAGIGDTPSLEDEAAGHVTQARMVSKSKDGTGSDDKKAKGADGKTKIATPRGAAPPGQKGQANATRIPAK**T**PPAPK**T**PPSSGEPPKSGDRSGYSSP 200

201 GSPG**T**PGSRSRTPSLPTPPTREPKKVAVVR**T**PPKSPSSAKSRLQTAPVPMPDLKNVKSKIGSTENLKHQPGGGKVQIINKKLDLSNVQSKCGSKDNIKHV 300

301 PGGGSVQIVYKPVDLSKVTSKCGSLGNIHHKPGGGQVEVKSEKLDFKDRVQSKIGSLDNITHVPGGGNKKIETHKLTFRENAKAKTDHGAEIVYK**S**PVV**S** 400

401 GDT**S**PRHLSNVSSTGSIDMVDSPQLATLADEVSASLAKQGL 441

**Fig. S13|** Sequence and information for native tau-441. The four domain sequences in native tau-441, along with theoretical isoelectric points (pI). The seven assumed potential phosphorylation sites for GSK3β-tau within this study are labelled (green circles). The illustration was made using IBS[29]. Amino acid residues are coloured according to their relevant domain, and assumed GSK3β phosphorylation sites are listed in bold.

#### SI-11 Resolved conformational dynamics of native tau-441 and GSK3β-tau

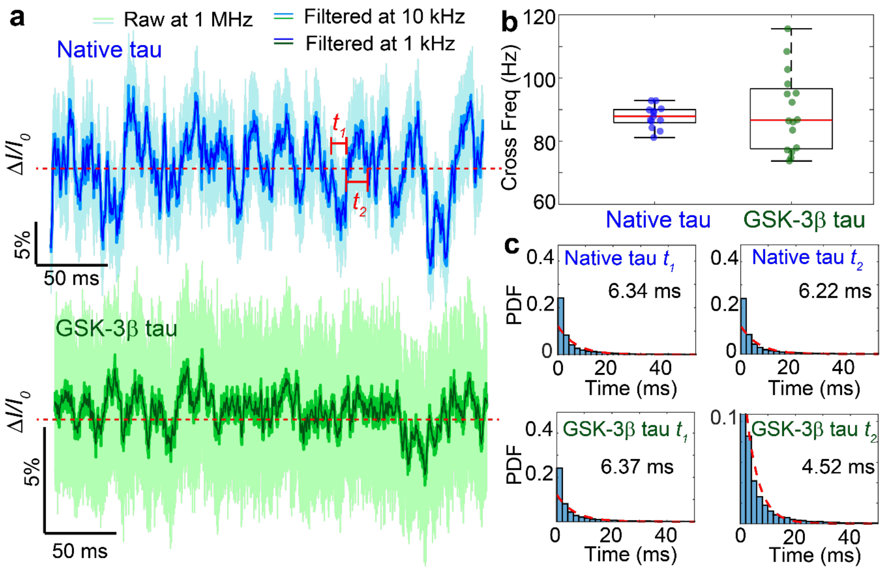


**Fig. S14|** Time scale of conformational dynamics of native tau-441 and GSK3β-tau. **a**, Zoomed traces of native tau-441 and GSK3β-tau, showing raw (1 MHz), and digital filtered (10 kHz and 1 kHz). No significant difference in dynamics was observed between 10 kHz and 1 kHz data, hence, the following dynamic analysis used 1 kHz-filtered data. The red dashed line represents the mean value of the signal. Time above ($t_1$) and below ($t_2$) the mean are defined as dwell times for fluctuation analysis. **b**, Frequency of mean-crossing events, reflecting conformational fluctuation rates. Both proteins exhibit a median crossing frequency of ~85 Hz, corresponding to a characteristic timescale of ~10 ms. c, Exponential fits to dwell time distributions above and below the mean. The observed ~6 ms average dwell time and exponential decay behaviour suggests the signal reflects the large-scale domain fluctuations in both IDPs.

### SI-12 Sequence information for Sam68 and G8.5 RNA

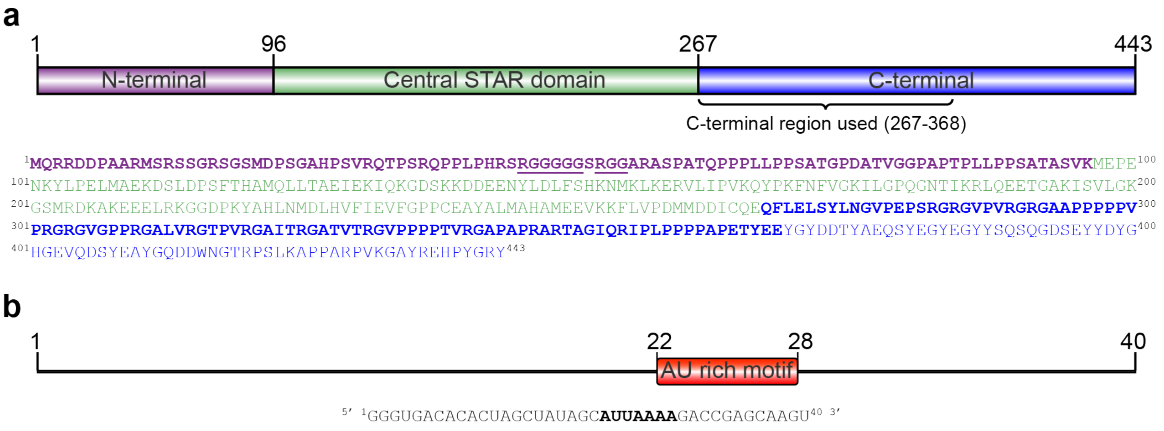


**Fig. S15|** Sequence information for Sam68 and G8.5 RNA. **a**, Sequence for Sam68 coloured by domain. Residues for the N- and C-terminal used in this study highlighted in bold[30]. Bold and underlined residues represent the RNA binding site–the RG rich region–on the N-terminus. **b**, Sequence for G8.5 RNA[31]. Bold letters represent the AU rich region involved in binding to the RG rich region on the N-terminal. Illustrations were made using IBS[29].

### SI-13 Trapping trajectory of the Sam68 N-terminal before and after detrend

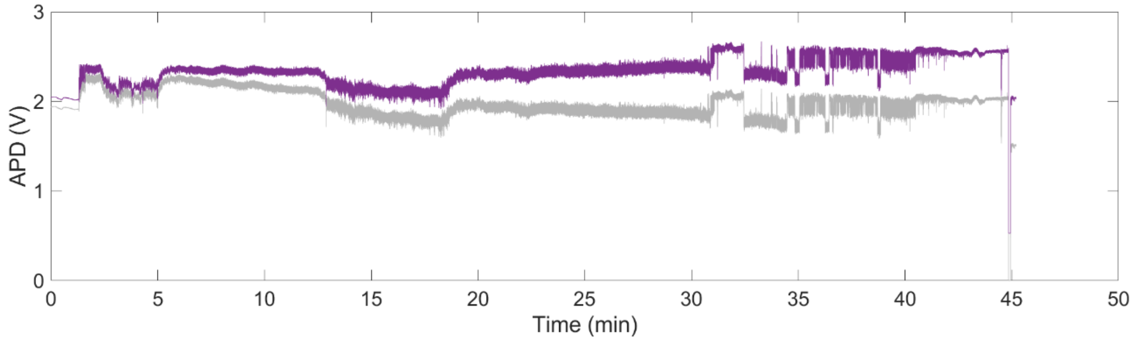


**Fig. S16|** Sam68 N-terminal trapping trajectory with and without detrend. Raw (grey) and detrended (purple) transmission trace of the Sam68 N-terminal shown in Fig. 4a. Data shown were filtered at 1 kHz.

### SI-14 Dissociation constant of the Sam68 N-terminal and G8.5 RNA binding interaction

To determine the dissociation constant ($K_d$) for the binding between the Sam68 N-terminal and G8.5 RNA (**Figs. 4e–g**), we analysed a ~7 min section of the trace where binding interactions occurred, indicated by the dashed box in **Fig. 4a**. The PDF on the right insert of **Fig. 4d** indicates four distinct levels in the binding trace. We assigned the lowest peak as only the Sam68 N-terminal, the highest peak to the fully bound Sam68 N-terminal–RNA complex (which stabilised into a rigid conformation afterwards), and the two intermediate states to transient configurations where both RNA and the Sam68 N-terminal coexisted in the trap.

We first applied step fitting to the 1kHz filtered trace using the *AutoStepfinder* package developed by Loeff and Kerssemakers[32]. We then generated a 3-step fit trace (blue trace in **Fig. 4e**) by assigning each point in the step-fit trace to the closest of three levels: the complex and two intermediate levels. The time constants were then extracted from the 3-step fit, defining binding time ($\tau_{\text{on}}$) as the duration of the middle and bottom levels (where the RNA is not fully bound), and dissociation time ($\tau_{\text{off}}$) as the duration of the top level (where RNA is fully bound). **Figures 4f** and **4g** show histograms of $\tau_{\text{on}}$ and $\tau_{\text{off}}$, overlaid with the exponential decay curves. These curves were obtained by fitting cumulative density functions (CDFs) of $\tau_{\text{on}}$ and $\tau_{\text{off}}$ using an integrated exponential decay function

$$f(CDF) = Bx - At \cdot e^{-\frac{x}{t}} + C \quad \text{(S8)}$$

with open parameters *A*, *B*, and *C*, and the time constant $t$ obtained from the fit. From this fit, we obtained the time constants $\tau_{\text{on}}$ = 1416 ms and $\tau_{\text{off}}$ = 177 ms.

The association rate constant ($k_{\text{on}}$) and dissociation rate constant ($k_{\text{off}}$) were then calculated using $\tau_{\text{on}}$ and $\tau_{\text{off}}$ obtained from **Eq. S8**, and the molar ligand concentration *C*[33–35]:

$$k_{\text{on}} = \frac{1}{C \cdot \tau_{\text{on}}} \quad \text{(S9)}$$

$$k_{\text{off}} = \frac{1}{\tau_{\text{off}}} \quad \text{(S10)}$$

With an RNA concentration of 1 μM, we calculated the rate constants as

$$k_{\text{on}} = \frac{1}{1.416[\text{s}] \times 10^{-6}[\text{M}]} = 7.062 \times 10^{5}\ [\text{M}^{-1}\text{s}^{-1}]$$

$$k_{\text{off}} = \frac{1}{0.177[\text{s}]} = 5.65\ \text{s}^{-1}$$

Therefore, the dissociation $K_d$ was determined to be 8 μM using the following relationship:

$$K_d = \frac{k_{\text{off}}}{k_{\text{on}}} \quad \text{(S11)}$$

#### SI-15 RNA binding to Sam68 N-terminal without dissociation

In some experiments, RNA binding transitioned the Sam68 N-terminal from an intrinsically disordered state to a more ordered complex without frequent binding/unbinding events (**Fig. S17a**). While such a trace cannot be used to determine the dissociation constant of the RNA binding interaction, we consistently observed increased changes in $\Delta I/I_0$ signal upon RNA binding (**Fig. S17b**), reflecting the greater hydrodynamic volume of the RNA-protein complex compared to the unbound Sam68 N-terminal. Meanwhile, the reduction in signal fluctuations (**Fig. S17c**) indicates the transition of the Sam68 N-terminal to a more compact, ordered structure, with dynamics occurring on timescales of hundreds of microseconds to seconds, as indicated in the PSD (**Fig. S17d**).

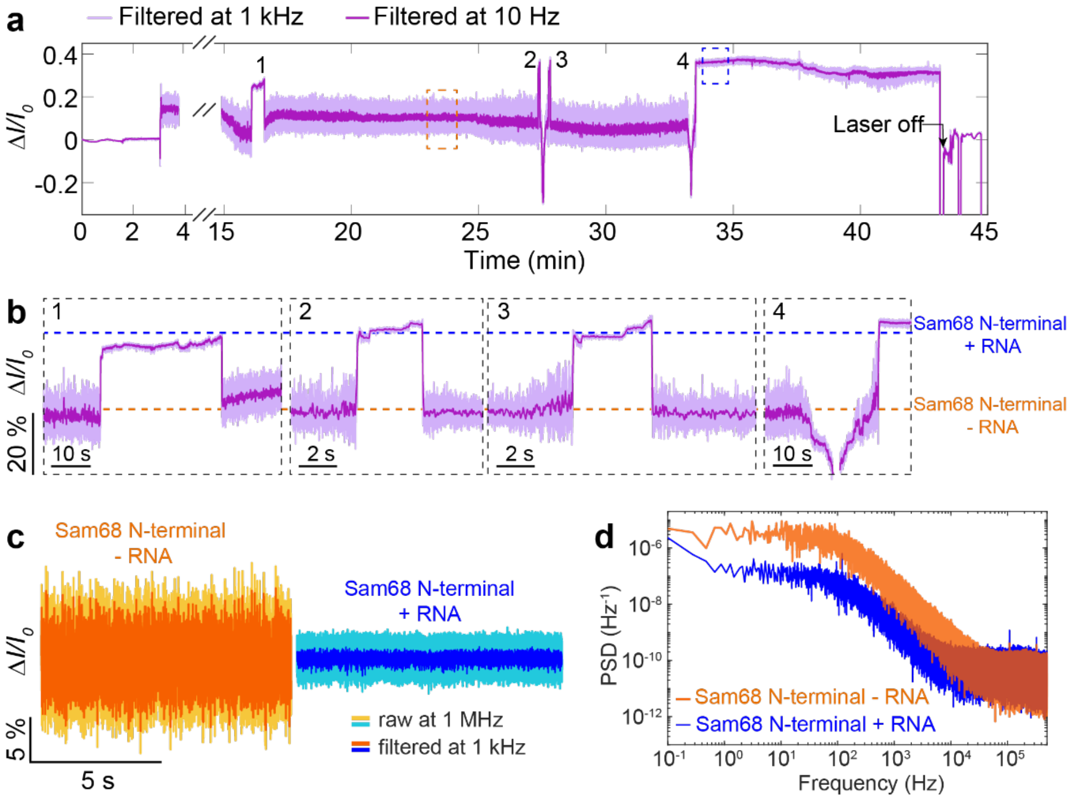


**Fig. S17|** Binding of G8.5 RNA to the Sam68 N-terminal. **a**, Trapping trace of Sam68 N-terminal followed by binding/unbinding of RNA. Data was filtered to 1 kHz (light purple) and 10 Hz (dark purple). **b**, Four binding/unbinding events depicting the reduction in signal fluctuation of the Sam68 N-terminal from unbound (orange) to RNA bound (blue) states. **c**, Zoomed segments from panel **a** depicting the Sam68 N-terminal before RNA binding (orange, from orange dashed box) and after RNA binding (blue, from blue dashed box). Traces are shown in raw data (1 MHz sampling rate, light colours) and filtered data (1 kHz, dark colours). **d**, PSD of normalised traces for the Sam68 N-terminal before RNA binding (orange) and after RNA binding (blue) for comparison.

### Supplementary references


1. Quinten Michael. Beyond Mie's Theory I – Nonspherical Particles. in *Optical Properties of Nanoparticle Systems* 255 (John Wiley & Sons, Ltd, 2011). doi:10.1002/9783527633135.ch9.
2. Pang, Y. & Gordon, R. Optical Trapping of a Single Protein. *Nano Lett.* **12**, 402–406 (2012).
3. Ying, C. *et al.* Watching Single Unmodified Enzymes at Work. Preprint at https://doi.org/10.48550/arXiv.2107.06407 (2021).
4. Peters, M. *et al.* Energy landscape of conformational changes for a single unmodified protein. *Npj Biosensing* **1**, 1–10 (2024).

5. Born, M. & Wolf, E. *Principles of Optics: Electromagnetic Theory of Propagation, Interference and Diffraction of Light*. (Cambridge University Press, Cambridge, 1999).
6. Li, L., Li, C., Zhang, Z. & Alexov, E. On the Dielectric “Constant” of Proteins: Smooth Dielectric Function for Macromolecular Modeling and Its Implementation in DelPhi. *J. Chem. Theory Comput.* **9**, 2126–2136 (2013).
7. Sarimov, R. M., Matveyeva, T. A. & Binhi, V. N. Laser interferometry of the hydrolytic changes in protein solutions: the refractive index and hydration shells. *J. Biol. Phys.* **44**, 345–360 (2018).
8. Booth, L. S. *et al.* Modelling of the dynamic polarizability of macromolecules for single-molecule optical biosensing. *Sci. Rep.* **12**, 1995 (2022).
9. Schuler, B. & Hofmann, H. Single-molecule spectroscopy of protein folding dynamics—expanding scope and timescales. *Curr. Opin. Struct. Biol.* **23**, 36–47 (2013).
10. Ying, C. *et al.* Formation of Single Nanopores with Diameters of 20–50 nm in Silicon Nitride Membranes Using Laser-Assisted Controlled Breakdown. *ACS Nano* **12**, 11458–11470 (2018).
11. Toral-Rios, D., Pichardo-Rojas, P. S., Alonso-Vanegas, M. & Campos-Peña, V. GSK3β and Tau Protein in Alzheimer’s Disease and Epilepsy. *Front. Cell. Neurosci.* **14**, (2020).
12. Alquezar, C., Arya, S. & Kao, A. W. Tau Post-translational Modifications: Dynamic Transformers of Tau Function, Degradation, and Aggregation. *Front. Neurol.* **11**, (2021).
13. Basheer, N. *et al.* Does modulation of tau hyperphosphorylation represent a reasonable therapeutic strategy for Alzheimer’s disease? From preclinical studies to the clinical trials. *Mol. Psychiatry* **28**, 2197–2214 (2023).
14. Beurel, E., Grieco, S. F. & Jope, R. S. Glycogen synthase kinase-3 (GSK3): Regulation, actions, and diseases. *Pharmacol. Ther.* **148**, 114–131 (2015).

15. Chatterjee, S., Sang, T.-K., Lawless, G. M. & Jackson, G. R. Dissociation of tau toxicity and phosphorylation: role of GSK-3β, MARK and Cdk5 in a Drosophila model. *Hum. Mol. Genet.* **18**, 164–177 (2009).

16. Darling, A. L. & Uversky, V. N. Intrinsic Disorder and Posttranslational Modifications: The Darker Side of the Biological Dark Matter. *Front. Genet.* **9**, (2018).

17. Marsh, J. A. & Forman-Kay, J. D. Sequence Determinants of Compaction in Intrinsically Disordered Proteins. *Biophys. J.* **98**, 2383–2390 (2010).

18. Jin, F. & Gräter, F. How multisite phosphorylation impacts the conformations of intrinsically disordered proteins. *PLOS Comput. Biol.* **17**, e1008939 (2021).

19. Lasorsa, A. *et al.* Magnetic resonance investigation of conformational responses of tau protein to specific phosphorylation. *Biophys. Chem.* **305**, 107155 (2024).

20. Jeganathan, S. *et al.* Proline-directed Pseudo-phosphorylation at AT8 and PHF1 Epitopes Induces a Compaction of the Paperclip Folding of Tau and Generates a Pathological (MC-1) Conformation *. *J. Biol. Chem.* **283**, 32066–32076 (2008).

21. Pearlman, S. M., Serber, Z. & Ferrell, J. E. A Mechanism for the Evolution of Phosphorylation Sites. *Cell* **147**, 934–946 (2011).

22. Chatterjee, S., Sang, T.-K., Lawless, G. M. & Jackson, G. R. Dissociation of tau toxicity and phosphorylation: role of GSK-3β, MARK and Cdk5 in a Drosophila model. *Hum. Mol. Genet.* **18**, 164–177 (2009).

23. Chakraborty, P. *et al.* GSK3β phosphorylation catalyzes the aggregation of tau into Alzheimer's disease-like filaments. *Proc. Natl. Acad. Sci.* **121**, e2414176121 (2024).

24. El Hajjar, L. *et al.* Effect of PHF-1 hyperphosphorylation on the seeding activity of C-terminal Tau fragments. *Sci. Rep.* **15**, 9975 (2025).

25. Lin, Y.-T. *et al.* The binding and phosphorylation of Thr231 is critical for Tau's hyperphosphorylation and functional regulation by glycogen synthase kinase 3β. *J. Neurochem.* **103**, 802–813 (2007).

26. Structural Polymorphism of 441-Residue Tau at Single Residue Resolution | PLOS Biology. https://journals.plos.org/plosbiology/article?id=10.1371/journal.pbio.1000034.

27. Adzhubei, A. A., Sternberg, M. J. E. & Makarov, A. A. Polyproline-II Helix in Proteins: Structure and Function. *J. Mol. Biol.* **425**, 2100–2132 (2013).

28. Chebrek, R., Leonard, S., de Brevern, A. G. & Gelly, J.-C. PolyprOnline: polyproline helix II and secondary structure assignment database. *Database* **2014**, bau102 (2014).

29. IBS 2.0: an upgraded illustrator for the visualization of biological sequences | Nucleic Acids Research | Oxford Academic. https://academic.oup.com/nar/article/50/W1/W420/6586865.

30. Malki, I. *et al.* Cdk1-mediated threonine phosphorylation of Sam68 modulates its RNA binding, alternative splicing activity and cellular functions. *Nucleic Acids Res.* **50**, 13045–13062 (2022).

31. Lin, Q., Taylor, S. J. & Shalloway, D. Specificity and Determinants of Sam68 RNA Binding: IMPLICATIONS FOR THE BIOLOGICAL FUNCTION OF K HOMOLOGY DOMAINS *. *J. Biol. Chem.* **272**, 27274–27280 (1997).

32. Loeff, L., Kerssemakers, J. W. J., Joo, C. & Dekker, C. AutoStepfinder: A fast and automated step detection method for single-molecule analysis. *Patterns* **2**, (2021).

33. Homola, J. Surface Plasmon Resonance Sensors for Detection of Chemical and Biological Species. *Chem. Rev.* **108**, 462–493 (2008).

34. Schuck, P. USE OF SURFACE PLASMON RESONANCE TO PROBE THE EQUILIBRIUM AND DYNAMIC ASPECTS OF INTERACTIONS BETWEEN BIOLOGICAL MACROMOLECULES1. *Annu. Rev. Biophys.* **26**, 541–566 (1997).

35. Zhang, P. *et al.* Label-Free Imaging of Single Proteins and Binding Kinetics Using Total Internal Reflection-Based Evanescent Scattering Microscopy. *Anal. Chem.* **94**, 10781–10787 (2022).
